# Studies of mice with a large deletion of the ARPKD-associated *Pkhd1* locus likely explain its GWAS association with glaucoma in humans

**DOI:** 10.64898/2026.02.15.706040

**Authors:** Yu Ishimoto, Luis F Menezes, Naoki Nakaya, Karla Barbosa, Yukihiro Horie, Teruhiko Yoshida, Jeff Reece, Fang Zhou, Stanislav Tomarev, Laura Kerosuo, Gregory G. Germino

## Abstract

*PKHD1*, the gene primarily mutated in human autosomal recessive polycystic kidney disease, is one of the top 20 genes associated with primary open angle glaucoma (POAG) and associated endophenotypes in Genome-Wide Association Studies. Here, we show that *Pkhd1^del3-67/del3-67^* mutant mice develop congenital glaucoma due to anterior segment dysgenesis. Using a combination of genetic, epigenetic, bioinformatics and mouse developmental biology approaches, we show that *Pkhd1^del3-67/del3-67^* mice lack *Tfap2b* and AP-2β expression in a subset of periocular mesenchymal cells at E13.5 and its derivatives. Our data suggest that the *Pkhd1^del3-67^* deletion disrupts features of the *Pkhd1-Tfap2b* genomic architecture essential for *Tfap2b* cell-specific activities. Consistent with this model, *Pkhd1^del3-67/+^;Tfap2b^ko/+^*trans-heterozygotes lack *Tfap2b* and AP-2β in relevant cell-types and have similar eye abnormalities as neural crest cell-specific *Tfap2b^ko^* mutants. These findings provide a likely causal explanation for how SNPs associated with *PKHD1* are functionally linked to POAG and add insight into understanding the complexity of disease-causing SNP associations and gene regulatory mechanisms.

## INTRODUCTION

Glaucoma is a multifactorial degenerative optic neuropathy and a leading cause of blindness worldwide. It has been estimated that 79.6 million people were affected by glaucoma globally in 2020 ^1^ and that number is expected to increase to 111.8 million by 2040 ^2^. Intraocular pressure (IOP) influences progression of glaucoma and currently is the only modifiable risk factor. The risk and subtypes of glaucoma vary among populations and countries, with a large genetic component to its etiology. Recent large multi-ethic Genome-Wide Association Studies (GWAS) studies have identified hundreds of genomic loci for primary open angle glaucoma (POAG) or related endophenotypes, high IOP and large vertical cup-to-disc ratio (VCDR) ^3–6^. While mutations in a small number of genes have been reported to cause autosomal dominant and recessive forms of the disease, most single nucleotide polymorphisms (SNP) have either an indirect, inferred connection to candidate genes or the link has been undefined ^5,6^. SNPs near the *TFAP2B/PKHD1* loci ^3–6^ on chromosome 6p12.2-12.3 are a notable example of the latter.

*TFAP2B* encodes AP-2β, a member of the AP-2 family of transcription factors that functions as both a transcriptional activator and repressor and is part of the conserved gene regulatory network that controls neural crest development ^7^. The neural crest is an embryonic stem cell population that forms from the ectodermal germ layer in the dorsal aspects of the developing neural tube during third week of gestation, from where they delaminate and migrate around the developing body and broadly contribute to organogenesis. These pleistopotent stem cells give rise to a myriad of cell types that resemble progeny of all germ layers, and neurocristopathies comprise a diverse group of congenital birth defects affecting an appreciable percentage of newborns ^8^. Neural crest cells (NCC) also play a significant role in eye development by giving rise to the periocular mesenchyme (POM), which contributes to the development of corneal endothelium and stroma, the stroma of the iris and ciliary body, and the trabecular meshwork ^9^.

As a member of the neural crest gene regulatory network, *TFAP2B* is expressed in pre-migratory NCC during their specification ^10^ and in migrating NCC and their developing derivatives within multiple organs and tissues across species ^10,11^. While these data might suggest a causal link between the GWAS data and *TFAP2B*, mutations in this gene in humans result in autosomal dominant Char syndrome. Affected individuals present with facial dysmorphism, cardiac defect patent ductus arteriosus and hand anomalies, but relevant eye findings have not been reported ^12^. Homozygous *Tfap2b* mouse mutants also have not been reported to have eye abnormalities, dying shortly after birth and presenting with a range of other phenotypes including renal cystic disease, congenital cardiac abnormalities, and failure of maintenance and differentiation of sympathetic neurons and their progenitors^13–15^. However, conditional inactivation of *Tfap2b* in cranial neural crest cells (NCC) in mice using a *Wnt1-Cre* recombinase results in anterior segment dysgenesis ^16^ and early onset glaucoma. Thus, *Tfap2b* may be a good candidate gene for POAG, but the gene maps approximately ∼650kb away from the SNPs linked to the disease, and in fact most reports and websites list *PKHD1* as the target gene^3–6^.

*PKHD1* is among the largest human genes, extending over 500kb, and its 67 core exons encode a 4074aa single transmembrane protein, fibrocystin^17–19^. The protein traffics to the primary cilium ^20^, undergoes Notch-like proteolytic processing, and has a short cytoplasmic C-terminus that, when released, traffics to the nucleus ^21^ and the mitochondrion ^22^. Both *PKHD1* and its murine ortholog undergo a complex and extensive pattern of alternative splicing, generating transcripts highly variable in size ^17,19^. The functional significance of this splicing pattern has been controversial ^23^.

The biological link between *PKHD1* and POAG is not immediately obvious. Mutations in *PKHD1* are the principal cause of human autosomal recessive polycystic kidney disease (ARPKD), which as its name implies, is an uncommon cystic disease of the kidney, affecting ∼1/20,000 livebirths. The kidney phenotype can be highly variable, ranging from massive *in utero* nephromegaly and oligohydramnios resulting in perinatal lethality to adult-onset kidney failure. Congenital hepatic fibrosis is a universal feature, and it often progresses to severe portal hypertension necessitating hepatic transplantation. Pancreatic cystic disease is an infrequent presentation ^24^. While ARPKD is considered a ciliopathy, and many pediatric-onset renal ciliopathies present with ocular manifestations ^25^, human ARPKD has not been associated with any eye abnormalities.

Numerous *Pkhd1* mutant mouse models have been generated to study human ARPKD, and while most faithfully reproduce the liver phenotype ^26^, the kidney phenotype is highly variable and never as severe as seen in humans. Although some models also present pancreatic cysts, none of the models develop eye abnormalities or other phenotypes not observed in humans. Because none of the mutations disrupted the entire gene, *Pkhd1*’s complex splicing pattern was considered a possible explanation for the attenuated renal presentation. To exclude this possibility, our group used a novel design to delete almost the entire *Pkhd1* genomic locus ^27^. Homozygous mutants lacked renal cysts, but 100% developed eye abnormalities with findings consistent with glaucoma, a surprising finding not previously associated with *Pkhd1* models ^23,28–34^.

The eye findings in mice with near-complete deletion of *Pkhd1* prompted further consideration of a possible role for *PKHD1* in eye development and function. There is, in fact, evidence to suggest a role for the primary cilium in development of the anterior segment of the eye. Conditional deletion of *Ift88*, a ciliary gene associated with an ARPKD-phenotype in mice, results in anterior chamber dysgenesis, a phenotype like that associated with conditional inactivation of *Tfap2b* ^35^. Loss of *Ift88* results in defective ciliary structure and function, potentially disrupting the function of fibrocystin.

Additional evidence suggesting a potential role for *Pkhd1* in the eye comes from studies of *Tfap2b*. The partial overlap of *Tfap2b* null mice phenotypes and ARPKD pointed to a possible functional connection between *TFAP2B* and the ARPKD gene ^13^. Subsequently, after the ARPKD gene had been identified as *PKHD1*, Wu *et al* reported that Ap-2b may be a common regulator of *Pkhd1* and *Cys1* (another ARPKD gene)^36^ and that *Tfap2b* deletion could result in kidney cystic disease by disrupting *Pkhd1* and *Cys1* expression. These data suggest that *Pkhd1* could also be a target of Ap-2b in the eye. Collectively, the published literature suggests that the GWAS associations with POAG in the chromosome 6p12.2-3 interval could point to differences in function of *PKHD1*, *TFAP2B*, or both as responsible for the phenotype.

Here, we utilized a series of mouse models, genetics, bioinformatics and developmental biology approaches to describe a likely causal explanation for how SNPs associated with the *PKHD1* locus are functionally linked to POAG and the transcription factor *TFAP2B*, which is essential for anterior chamber development in the eye.

## RESULTS

### Genetic and functional genomic information related to *PKHD1/TFAP2B*

We started by analyzing the loci that are associated with POAG and IOP. The results show that *PKHD1* is one of the top 15 genes associated with the phenotype (Figure 1a) ^37^ Chromosome 6p12.2-3 SNPs with the highest association with POAG/IOP cluster near exon 67, just outside of the *PKHD1* gene itself, and are associated with lower risk (Figure 1b, c). SNPs with lower but still significant associations map along the length of *PKHD1* and the neighboring gene *TFAP2B* (Figure 1b, c). Interestingly, the SNP with the highest association to POAG that maps near the 3’ end of *PKHD1* is predicted to affect a binding site for *Msx2*, a transcription factor essential for normal eye development ^38^. Figure 1d presents the list of *Pkhd1* mutant mouse models that have been reported and the exonic location of their mutations, also graphically illustrated in Figure 1b (bottom). The *Pkhd1^del3–67^* mouse line is the only one that has the entire genomic interval deleted.

**Fig. 1:**
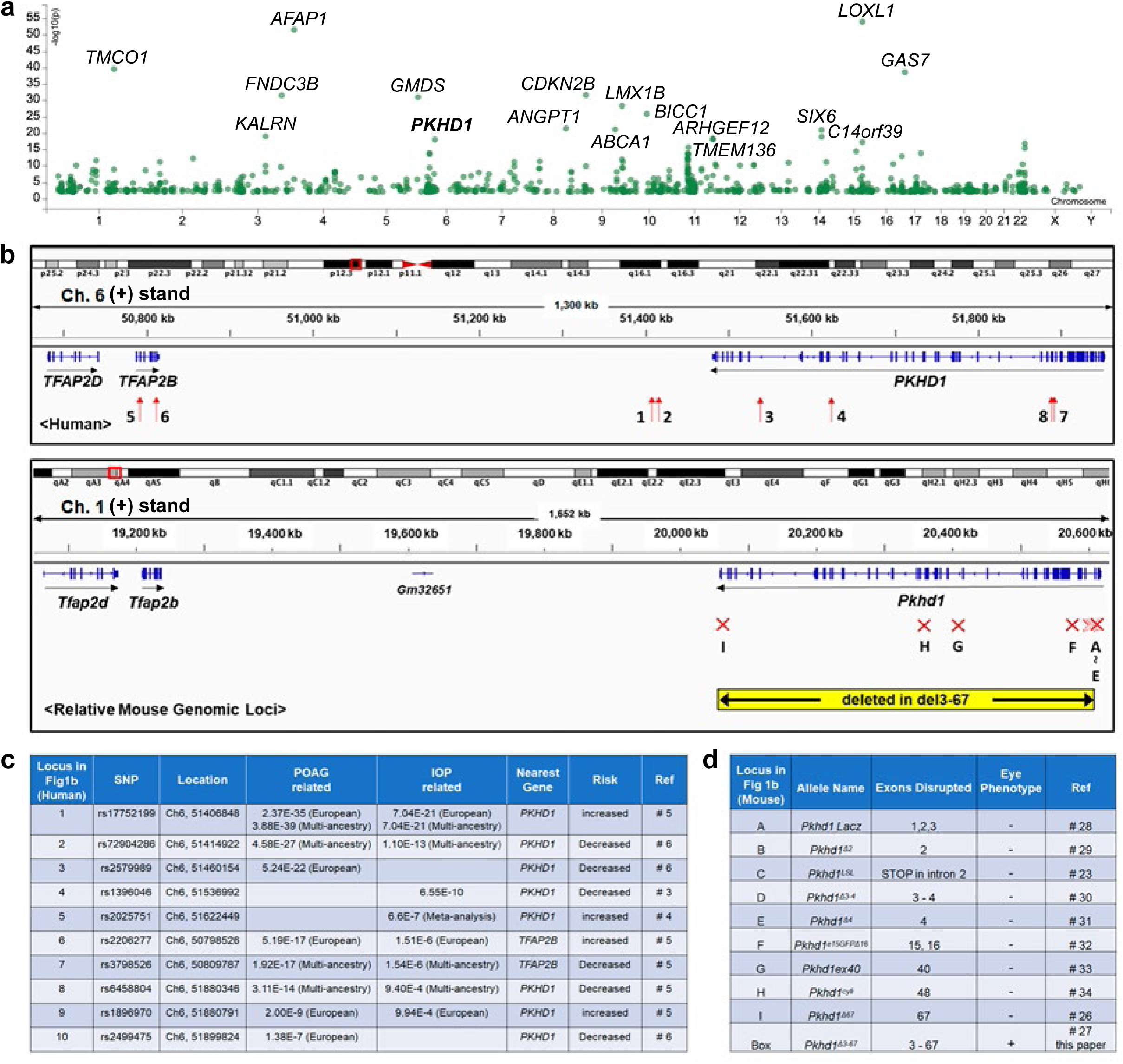
Genetic and functional genomic information related to the *PKHD1/TFAP2B*. (a) Loci associated with Primary Open Angle Glaucoma (POAG). The x-axis represents the chromosomal location of the loci and y-axis shows the -log10(p-value) of the association. Genes with association p < 1.1E-18 are labeled. (b) (Top) Genomic map of human *PKHD1* and *TFAP2B* loci based on the NC_000006.11 Chromosome 6 Reference GRCh37.p13 Primary Assembly visualized with Integrative Genomics Viewer (IGV). Red arrows in the box immediately below the map indicate approximate position of SNPs shown in Figure 1c. The numbers next to each arrow indicate the SNP identified in Figure 1c. (Bottom) Genomic map of mouse *Pkhd1* and *Tfap2* loci based on the Chromosome 1 Reference GRCm39/mm10 Primary Assembly visualized with IGV. Red “X”s show the relative locations of mutations introduced into the mouse orthologue. The letter below each “X” identifies the mouse model described in Figure 1d. Note that the corresponding mouse and human genomic regions are very similarly structured. (c) Previously reported SNPs in *PKHD1/TFAP2B* genomic loci associated with POAG and intraocular pressure (IOP). SNP location was taken from the NC_000006.11 Chromosome 6 Reference GRCh37.p13 Primary Assembly. (d) Previously reported Pkhd1 mutant mouse models. Letters listed in the first column identify the genomic position of their mutation in Figure 1b.

### *Pkhd1^del3-^*^67*/del3-*67^ develop early-onset glaucoma

We recently reported ^27^ that in four distinct mouse lines derived from independently derived founder mice in which *Pkhd1* exons 3 to 67 were deleted by Cre-recombination, all homozygotes with deletion of *Pkhd1* exons 3-67 (*Pkhd1^del3-67/del3-67^*) develop corneal opacities in both eyes detectable within the first 3-4 weeks of life (Supplementary Figure 1a). Their eye globes become enlarged as compared to controls, and this eye phenotype is specific for *Pkhd1^del3-67/del3-67^* as none of the *Pkhd1^del3-67/+^*heterozygotes develop it (Supplementary Figure 1b, c). We found no change in the size of eye globes normalized for body weight at 1 week of postnatal age (Supplementary Figure 1d), but they were significantly larger at 4 weeks of age though with some variability as they further aged (Supplementary Figure 1e, f). We observed no differences at any time point based on sex (Supplementary Figure 1g-i). These results suggest that eye abnormalities occur early in life before they become adult but are progressive in nature.

Since bulging eyes and eye opacities are often associated with severe glaucoma in humans, we evaluated IOP at three weeks of age, which is soon after mice open their eyes. As expected, IOP was significantly elevated in *Pkhd1^del3-67/del3-67^* pups compared with that of controls (Figure 2a). In humans, IOP measured by rebound tonometry correlates with corneal membrane thickness (CMT) so we determined CMT in P21 control and *Pkhd1^del3-67/del3-67^* eyes (Figure 2b, c) and found that it was moderately higher in *Pkhd1^del3-67/del3-67^* mice. The increase in IOP is out of proportion to the change in CMT, suggesting that IOP is truly elevated, but we cannot easily determine how much the increased CMT is contributing to this.

**Fig. 2:**
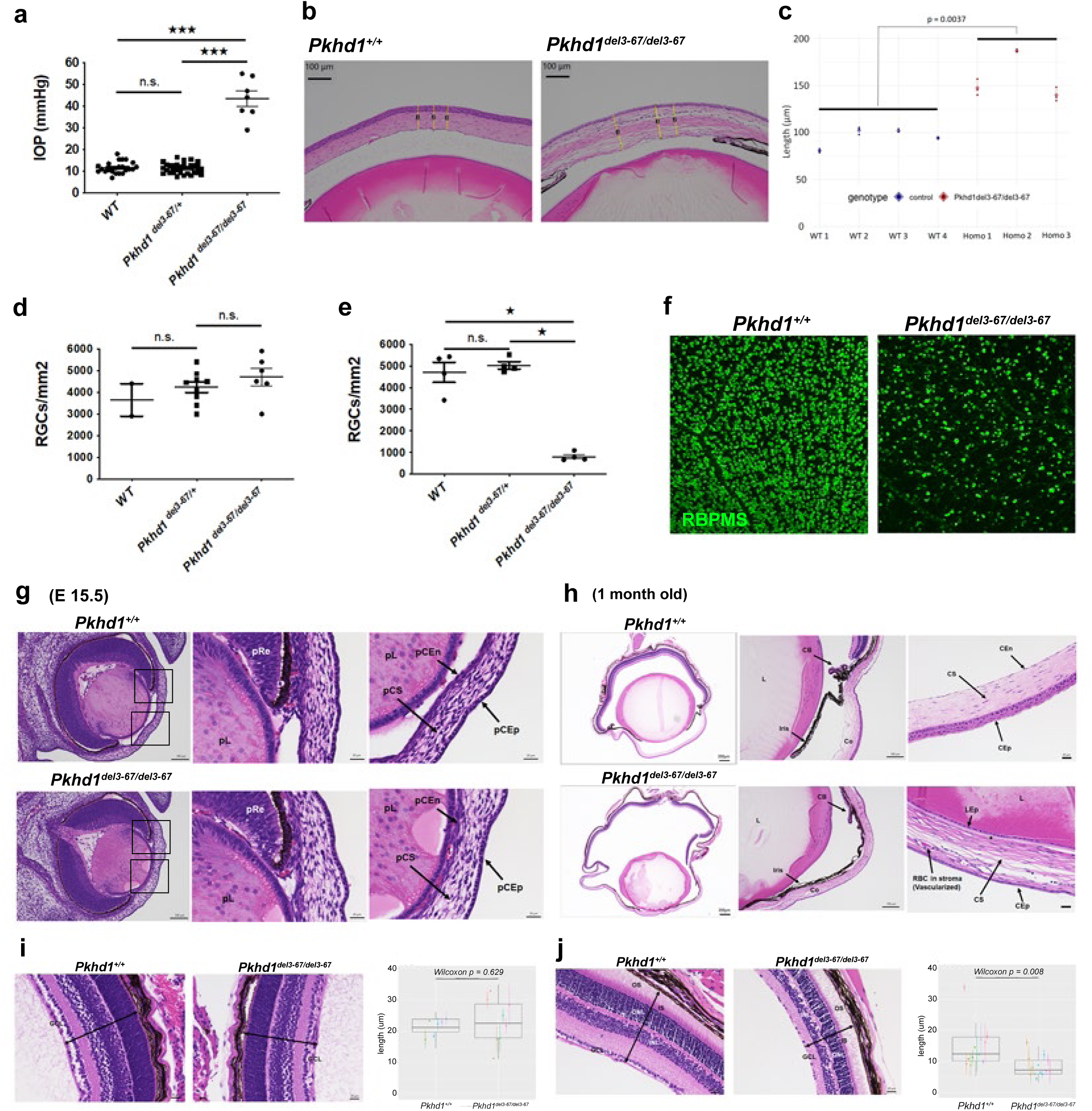
Ocular phenotype of *Pkhd1^del3-67/del3-67^*. (a) IOP of *Pkhd1^del3-67^* mouse line at 3 weeks of age. Each dot indicates IOP of one mouse (mean of right and left eye IOP measurements). ^★★★^ P < 0.001. n.s.. not significant. (b) Representative images of cornea at 3 weeks of age. Yellow lines indicate measurements of corneal thickness. (c) Corneal thickness of *Pkhd1^+/+^* (WT) (N=4) and *Pkhd1^del3-67/del3-67^* (Homo) (N=3) at 3 weeks of age. Each dot represents one of three measurements from a single eye of each mouse, and the mean values of the measurements from each mouse were compared. (d, e) Number of RGC at 1 (d) and 4 (e) weeks of age. Each dot represents the mean number of RGC counted from a series of immunostained images from a single mouse. ^★^ P < 0.05. n.s.. not significant. (f) Representative images of RGC at 4 weeks of age stained for RBPMS. Scale bar 100μm. (g, h) Representative eye pathology with H&E staining. Left panels: whole eye image, scale bar 100 μm (g) and 200 μm (h). Black squares indicate the region of angle tissue and presumptive cornea (g). Middle panels): angle tissue, scale bar 20 μm (f) and 100 μm (g). Presumptive cornea, scale bar 20 μm in both “g” and “h”. pL: presumptive lens, pRe: presumptive retina, pCS: presumptive corneal stroma, pCEn: presumptive corneal endothelium, pCEp: presumptive corneal epithelium. CB: ciliary body, Co: cornea, CEn: corneal endothelium, CS: corneal stroma, CEp: corneal epithelium, L: lens, LEp: lens epithelium. The star indicates absent corneal endothelium in *Pkhd1^del3-67/del3-67^*. (i, j) H&E-stained retinas and box plots showing ganglion cell layer thickness in *Pkhd1^del3-67/del3-67^* (N=4, 7) and *Pkhd1^+/+^* (N=3, 6) at 1 week (i) and 1 month (j). Each dot is the average thickness and the range indicates maximum and minimum values per eye; same color dots identify left/right eye values of the same animal. Scale Bar 20 μm. OS: outer segment, IS: inner segment, ONL: outer nuclear layer, INL: inner nuclear layer, GCL: ganglion cell layer. Two-headed arrows indicate thickness of retina.

While an elevated IOP is a major risk factor for glaucoma, our elevated values also could be the result of the abnormal cornea structure, so we looked for other evidence of glaucoma. Loss of retinal ganglion cells (RGC) is another prominent feature of glaucoma so we evaluated the number of RGC at one week and four weeks of age (Figure 2d, e). We found no difference in the RGC count in one week old *Pkhd1^del3-67/del3-67^* mutant pups compared with controls (Figure 2d), but it was significantly decreased at four weeks of age (Figure 2e, f). These results indicate that *Pkhd1^del3-67/del3-67^* mice develop manifestations of early-onset glaucoma with RGC damage that likely results from an elevated IOP.

### *Pkhd1^del3-67/del3-67^* embryos develop anterior segment dysgenesis

Retinal structure can be identified at E13.5 ^39^ and becomes fully functionally mature at around one month of age. To better define the basis for the elevated IOP and retinal damage, we evaluated the histopathology of *Pkhd1^del3-67/del3-67^* eyes from embryonic, one week and one month old mice, respectively. There were no obvious abnormalities in *Pkhd1^del3-67/del3-67^* mice at E13.5 (Supplementary Figure 2a). In contrast, by E15.5 when the corneal endothelium and the anterior chamber are normally fully formed in the control littermates, the cornea was attached to the lens in *Pkhd1^del3-67/del3-67^* mice (Figure 2g). At one week of age, the irido-corneal angle was closed with the lens attached to the cornea (Supplementary Fig 2b, c). At this younger age, the appearance of the corneal epithelium and endothelium of *Pkhd1 ^del3-67/del3-67^* mice were somewhat variable, being slightly flattened in most specimens, and red blood cells were visible in the stroma. The changes at one month were more pronounced. The anterior chamber of *Pkhd1^del3-67/del3-67^* mice was flattened or absent, with a rudimentary ciliary body (Figure 2h, Supplementary Figure 2d). The corneal endothelial layer of *Pkhd1^del3-67/del3-67^* mutants was mostly absent and the stroma was vascularized. The changes in the corneal epithelial layer were striking, with stratification greatly reduced or lacking in all mutants and keratitis in a subset (Supplementary Figure 2e). In sum, our results show that the *Pkhd1^del3-67/del3-67^* mice present with a severe developmental defect of the anterior segment that onsets by embryonic day E15.5.

Rod and cone photoreceptors have primary cilia, and retinal degeneration is a highly penetrant phenotype in ciliopathies ^40^. We therefore next evaluated the thickness of each layer of the retina at one week and one month of age. In one week old pups, no statistically significant evidence of abnormal morphology or thinning of any layers was found in *Pkhd1^del3-67/del3-67^* mice as compared to controls as visualized and quantified by using histochemistry (Figure 2i) whereas at one month of age, *Pkhd1^del3-67/del3-67^* mice had a thinned RGC layer (Figure 2j) and a trend toward a lower overall thickness of the entire retina. While the width of several layers was statistically different at a p-value of 0.05, the thickness of the rod and cone layers was nearly identical and consistent with preserved rod and cone structure (Supplementary Figure 3).

These results suggest that the eye abnormality in *Pkhd1^del3-67/del3-67^* starts in later embryonic stages and progresses to angle closure after birth, which causes elevated IOP and secondary retinal damage. Taken together, these findings are consistent with anterior segment dysgenesis and congenital glaucoma.

### *Pkhd1* expression is detectable in adult, but not in the developing, eye

Median tissue-specific *PKHD1* transcript expression is highest in the eye when searched using the Common Metabolic Diseases Knowledge Portal (Supplementary Figure 4a) ^41^, suggesting a possible novel function for *PKHD1* in eye development or function. However, none of the previously described mouse models, which had truncating mutations distributed along much of the length of *Pkhd1,* had an eye phenotype (Figure 1b, d). This prompted us to search for either an alternative eye-specific *Pkhd1* transcript or another gene embedded within the *Pkhd1* locus ^27^. For this, we queried publicly available single-nuclei RNA-seq (snRNA-seq) databases of human adult ocular anterior segment cell types ^42^ and found *PKHD1* was expressed widely at a very low level throughout the eye except in corneal endothelial cells, where it was expressed at higher levels (Supplementary Figure 4b, c, e, f).

Corneal endothelial, stromal and angle tissues, which are abnormal in *Pkhd1^del3-67/del3-67^* mice, are all derived from cranial neural crest cells (NCC) ^9,43^. *Tfap2b* is an important factor in early NCCs and their derivatives that contribute to eye development and its selective loss in NCC results in a phenotype like that of *Pkhd1^del3-67/del3-67^* mice ^16^. Therefore, we asked whether there is a functional connection between these two genes in the ocular anterior segment and found that *TFAP2B* expression was co-expressed with *PKHD1* in human corneal endothelial cell clusters (Supplementary Figure 4d, g). To determine if the pattern was conserved, we analyzed additional publicly available scRNA transcriptomes of adult mouse irises and corneas^44,45^ and found *Pkhd1* broadly expressed at very low levels, but with higher expression in a stromal population (“stroma 2”) thought to be derived from a subset of migratory NCC that also had high levels of *Tfap2b* expression (Supplementary Figure 4h-j). Next, we looked for evidence of an eye-specific transcript. For this, we determined *Pkhd1* expression for each exon and indeed found a disproportionally higher number of reads for exons 54 and 55, most notably in the NCC-derived stroma 2 population (Supplementary Figure 5a).

These data raised the possibility that *Tfap2b* could regulate a *Pkhd1* alternative transcript(s) in cells that form the anterior segment of the eye. There is precedence for this hypothesis: as indicated in the Introduction, a prior study reported that *Tfap2b* regulates *Pkhd1* expression in the kidney, thus explaining why *Tfap2b* mutants develop cystic kidney disease ^36^. While we identified AP-2β enhancer binding sites near exons 54 and 55 (Supplementary Figure 5b), RNAscope probe sets for full length *Pkhd1* and just for a short isoform that includes exons 54-56 failed to produce specific signals above background, indicating that expression levels were below detection for both (Supplementary Figure 6). Collectively, out data suggest that the abnormalities in eye development of *Pkhd1^del3-67/del3-67^* mice are unlikely due to loss of function of the *Pkhd1* protein.

### AP-2β and *Tfap2b*-expression are markedly decreased in *Pkhd1^del3-67^* mutant eyes

Given the similarity in eye phenotype of *Tfap2b^cko/cko^;Wnt1-Cre* and *Pkhd1^del3-67/del3-67^* mutant mice, we examined the pattern of AP-2β expression in the eyes of *Pkhd1^del3-67/del3-67^* mice. Starting at 1 month of age when there were obvious morphologic abnormalities, our results show that AP-2β and ZO1, the latter a marker for corneal endothelial cell integrity ^46^, were absent from where the corneal endothelium of *Pkhd1^del3-67/del3-67^* should have been (Figure 3a, b; Supplementary Figure 7a, b). These results are consistent with prior reports that indicate *Tfap2b* is important in corneal endothelial cell development ^47^. We next analyzed at E15.5, a developmental stage with clear histopathologic abnormalities (Fig 2g), and confirmed the absence of AP-2β in corneal endothelium and stroma (Figure 3c, Supplementary Figure 7c). Analysis at an earlier time point (E13.5), which did not present any obvious morphological changes in the eyes of *Pkhd1^del3-67/del3-67^* mice (Supplementary Figure 2a), showed that AP-2β expression was already absent from the NCC-derived mesenchyme between the lens and surface ectoderm that gives rise to the future corneal endothelium and stroma (Figure 3d). However, AP-2β expression was still present in some cells of the periocular mesenchyme, which is derived from both the mesoderm and cranial neural crest (Supplementary Figure 7d) ^48^. We confirmed that the loss of AP-2β expression was due to decreased expression of the *Tfap2b* transcript in the same cell types (Figure 3e).

**Fig. 3:**
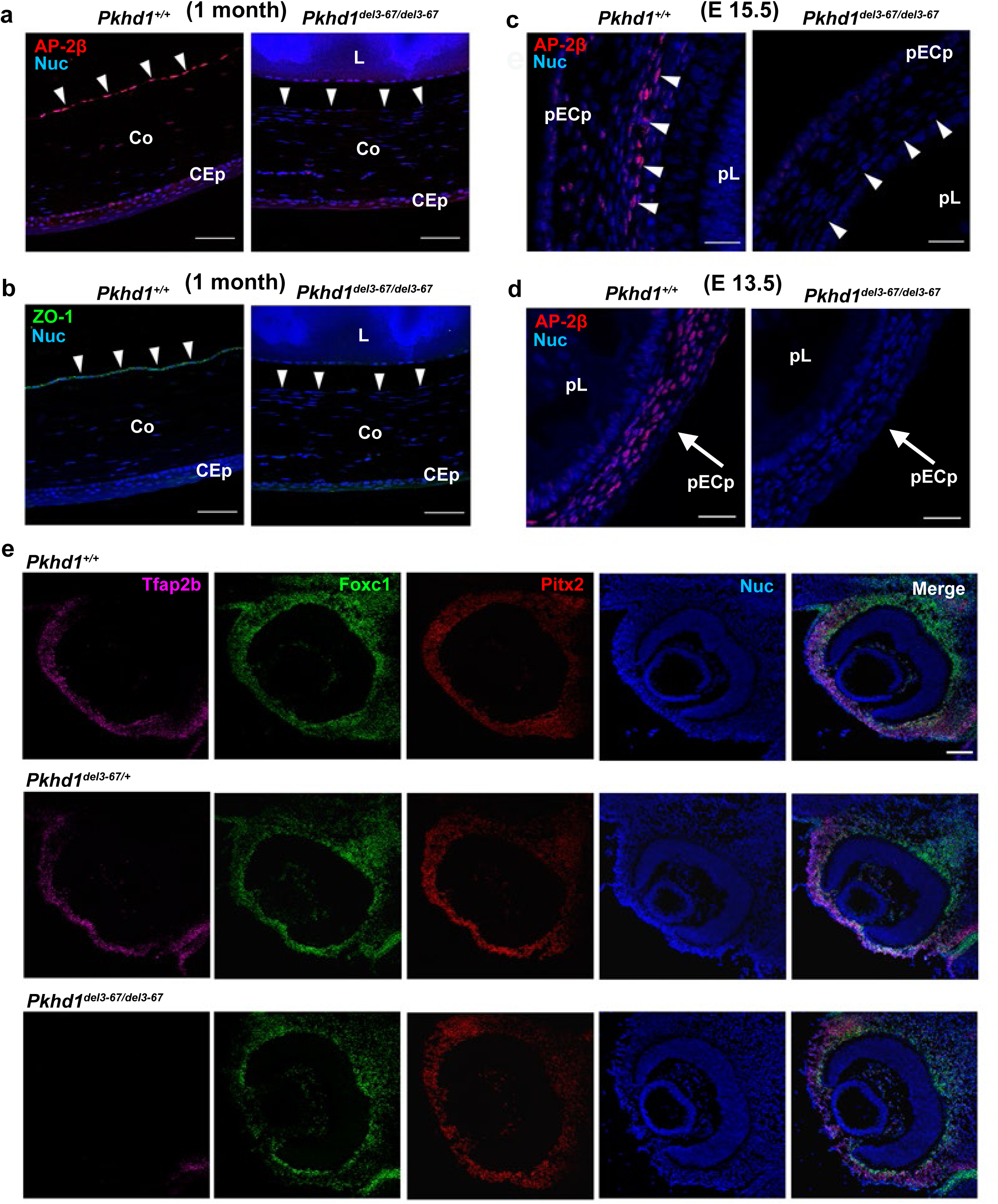
AP-2β and *Tfap2b* expression in adult and fetal control and *Pkhd1^del3-67/del3-67^* mouse eyes. (a, b) One month old littermate mouse eyes immunostained for AP-2β (a), ZO-1 (b) and Hoechst 33342. Scale bar 50 μm. Arrowheads indicate corneal endothelial cells. CEp: corneal epithelium, Co: cornea, L: lens. (c, d) Fetal littermate mouse eyes stained for AP-2β. Scale bar 25 μm in both panels. pCEp: presumptive corneal epithelium, pL: presumptive lens. (E15.5; n = 6 eyes from 3 mice for each group, E13.5; n = 12 eyes from 6 mice for each group). (e) Representative images of *in situ* hybridization of *Tfap2b*, *Foxc1*, and *Pitx2* in E13.5 eyes using RNAScope HiPlex v2 and stained with Hoechst 33342. Scale bar 50 μm.

Since we detected a loss of AP-2β expression in neural crest derivatives in the eye, and since AP-2β is broadly expressed in all NCC at the pre-migratory and migratory stages before they populate various target tissues, we asked whether NCC development was affected already during earlier embryonic stages in *Pkhd1^del3-67/del3-67^* mice. As neural crest cells are specified in the dorsal aspects of the neural tube, a cascade of transcription factors, including Sox9 and Sox10, are activated at different stages of the maturation process. Sox9 is expressed in the early pre-migratory NCCs whereas Sox10 is first activated when NCCs go through epithelial-mesenchymal transition to delaminate from the neural epithelium. Sox10 can therefore be used as a marker for newly as well as fully migrating neural crest cells that enter different target organs. To fully cover both stages of neural crest development, we used these two markers accordingly to understand whether the neural crest at these pre-migratory and migratory stages was impacted in *Pkhd1^del3-67/del3-67^* mice.

We examined the patterns of expression of AP-2β and Sox9 at E8 and found similar patterns of expression in pre-migratory NCCs in *Pkhd1^del3-67/del3-67^* mice and WT littermates (Figure 4a). Similarly, a day later at E9, when Sox10-positive NCCs have reached their target organs, including the eye cup, we found that AP-2β -positive NCCs had normally populated the eye in both WT and *Pkhd1^del3-67/del3-67^* mice (Figure 4b-h’). Additionally, the results showed AP-2β- positive NCCs were normally populating all other destination organs in *Pkhd1^del3-67/del3-67^* mice (Supplementary Figure 8). Collectively, these results suggest that *Pkhd1 ^del3-67/del3-67^* mice have normal neural crest development and *AP-2*β expression throughout the migrating stage, and that the NCC normally reach their target organs. However, once in the eye cup, the NCC-derived POM region no longer expresses *AP-2*β normally.

**Fig. 4:**
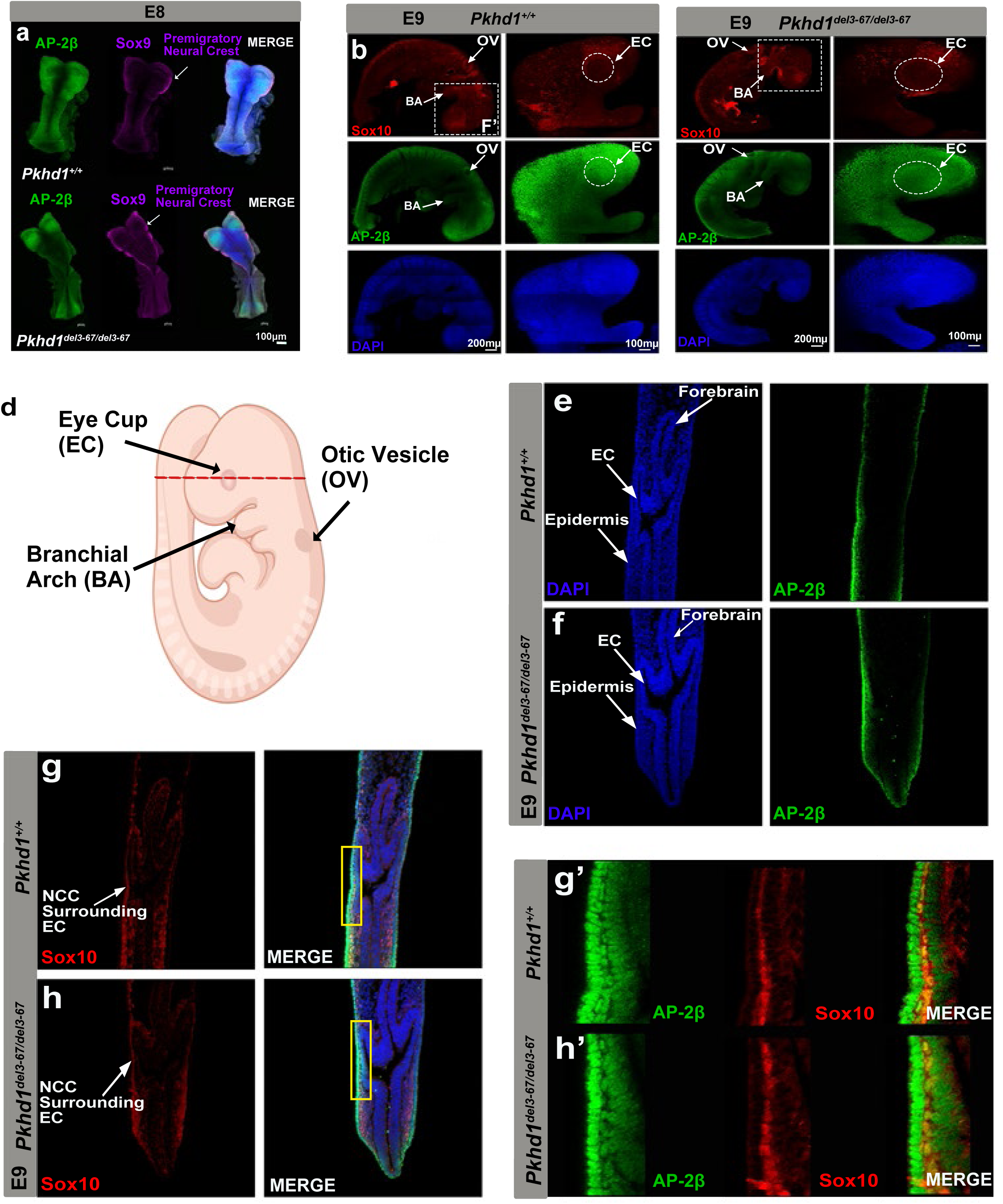
AP-2β, Sox 9 and Sox 10 expression in *Pkhd1^+/+^* control and *Pkhd1^del3-67/del3-67^* mouse embryos. (a) Whole mount immunostaining of E8 mouse embryos stained for AP-2β, Sox9, and DAPI (blue). (b, c) Whole mount E9 mouse wild type and *Pkhd1^del3-67/del3-67^* embryos stained for AP-2β (green), Sox10 (red), and DAPI (blue). Insets present a magnification of the head region. (d) Schematic diagram of an E9 mouse embryo depicting with a red line the respective plane of the cross section presented in Figure 4e-h. (e-h) Cryosections from the forebrain level stained for AP-2β and Sox10. Arrowheads point to NCCs surrounding the eye cup. (g’, h’) Yellow insets show a magnification of the squared area in Figure 4g, h showing an overlap of AP-2β and Sox10 in the neural crest and its derivatives.

Finally, to exclude the small possibility that the Cre-based method used to generate the *Pkhd1^del3-67^*allele inadvertently caused a chromosome rearrangement or deletion affecting the *Tfap2b* locus, we re-analyzed the whole genome sequence files that were previously reported in the initial characterization of the *Pkhd1^del3-67^* mouse line^27^ and confirmed that the *Tfap2b* locus was unaffected (Supplementary Figure 9).

### Tfap2b^ko/+^; Pkhd1^del3-67/+^ trans-heterozygous mice

The observation that *Tfap2b* expression is greatly reduced in *Pkhd1^del3-67/del3-67^* mice suggests that deletion of *Pkhd1* disrupts overall local chromosome architecture and indirectly effects *Tfap2b* expression. Strong evidence in support of this model can be found in public databases of genomic interaction domains. Results from published human and mouse datasets suggest that *PKHD1* and *TFAP2B* are part of the same Topologically Associated Domain (TAD), showing evidence of chromatin interaction between different regions of this genomic interval in various tissues (Figure 5). These data suggest that the effects of the *Pkhd1^del3-67^* deletion would be exerted in “cis” as local chromatin structure would control allelic expression. Thus, it can be expected that the eye phenotype of trans-heterozygous mice carrying one null allele for *Tfap2b* and the other with the large *Pkhd1* deletion would functionally look like *Tfap2b^cko/cko^;Wnt1-Cre* or *Pkhd1^del3-^*^67/del3-67^ mice.

**Fig. 5:**
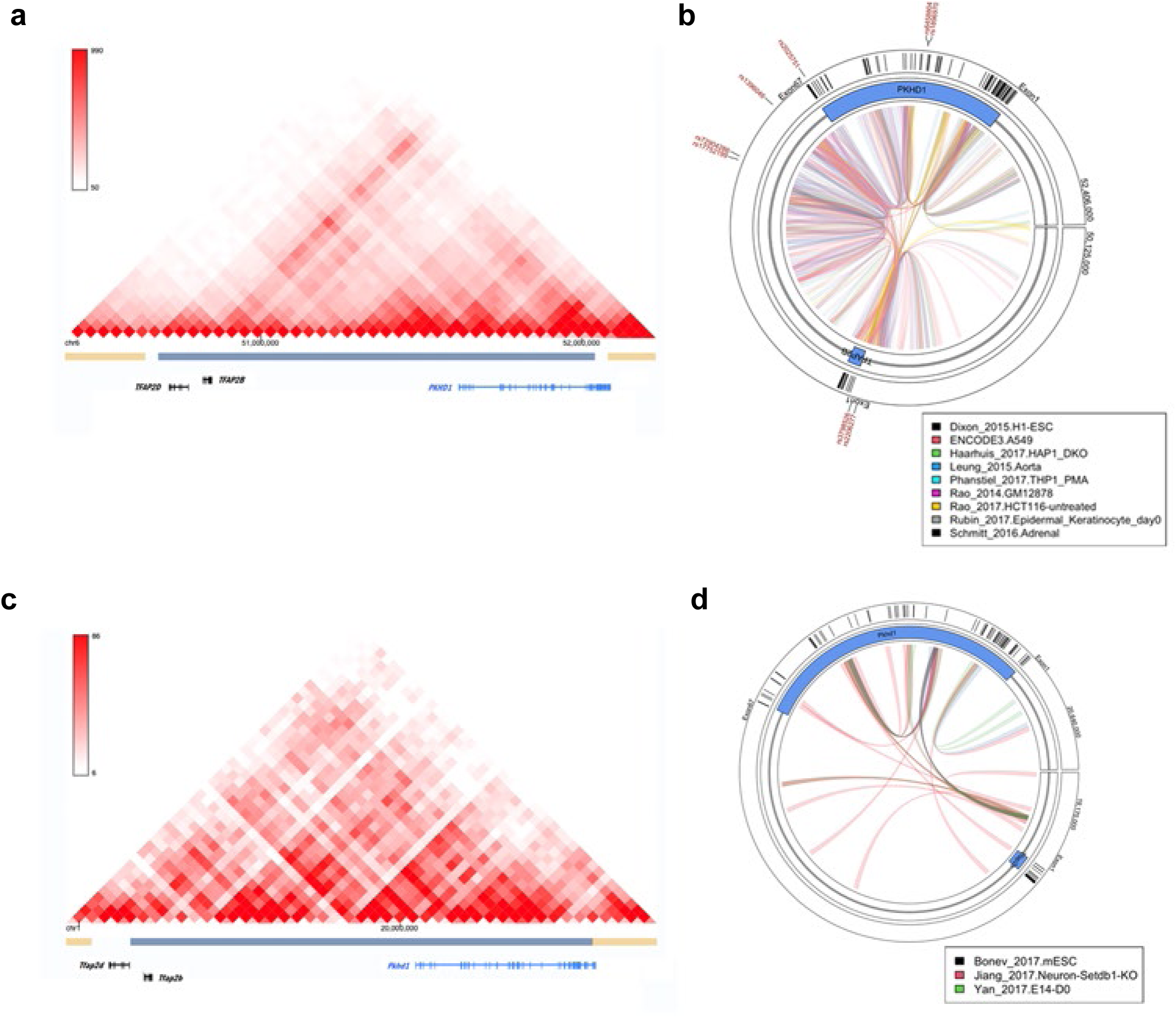
Topologically associated domains in human and mouse genome. (a, c) Results from published datasets suggest that human and mouse *PKHD1* and *TFAP2B* are part of the same Topologically Associated Domain (TAD). (b, d) Chromatin interaction between different regions of the *PKHD1-TFAP2B* loci in various tissues. Chromatin links are colored by publication in which they were reported. The approximate location of human SNPs associated with glaucoma is shown in the outer rim of panel b.

To test this hypothesis, we deliberately used *Tfap2b* null mice (*Tfap2b^ko/ko^*) to control for possible variability in Cre activity. We also wanted to exclude possible confounding effects resulting from use of the Wnt1-Cre. It has been reported to cause ectopic activation of Wnt signaling and was used to conditionally inactivate *Tfap2b* and *Ift88* in NCCs ^16,35,49^. Given that the *Tfap2b* and *Ift88* conditional mutants had similar phenotypes and both genes regulate Wnt signaling, we could not exclude ectopic Wnt signaling as a common contributor to their shared phenotypes.

In parallel with conducting crosses between *Pkhd1* and *Tfap2b*, we also sought to evaluate the eyes of *Tfap2b* homozygous null mice as previous reports ^50^ had not described any eye abnormalities. As previously reported for *Tfap2b^ko/+^* x *Tfap2b^ko/+^* crosses, all genotypes were present at E15.5 at expected Mendelian ratios, but the ratios were markedly distorted by the time of weaning with many mice lost in the perinatal period (Supplementary Figure 10a-f). The sole *Tfap2b^ko/ko^* survivor had mild bilateral renal cystic disease, as had been previously reported (Supplementary Figure 10g, h), but also had developed gross eye abnormalities like those described for *Pkhd1^del3-67/del3-67^* mice (Supplementary Figure 11a). Histopathological evaluation confirmed the two mutant lines had similar pathology (Supplementary Figure 11b, c). *Tfap2b^ko/ko^* did not, however, develop cystic liver disease like that of *Pkhd1^del3-67/del3-67^* mice (Supplementary Figure 10i). Because there was only one liveborn available for analysis, we examined the eyes of *Tfap2b^ko/ko^* embryos at E15.5 and confirmed that the phenotype was consistently present (Supplementary Figure 11d).

For the *Pkhd1^del3-67/+^*and *Tfap2b^ko/+^* crosses, we evaluated 97 pups from 12 litters and found all genotypes born at the expected frequency without perinatal lethality (Figure 6a). As expected, all trans-heterozygous mice developed eye abnormalities like those of *Pkhd1^del3-67/del3-67^* mutants (Figure 6b-d). Their eyes were generally enlarged with high IOP (Figure 6e, f, Supplementary Figure 12a-c) and the number of retinal ganglion cells (RGC) was decreased at the age of four weeks (Figure 6g, h, Supplementary Figure 12d). The *Tfap2b^ko/+^; Pkhd1^del3-67/+^* trans-heterozygous mice eyes showed similar histopathology with corneal, stromal and endothelial abnormalities, iris attachment, closed angles (Figure 6i), although quantification of their respective retinal layers suggested that the *Tfap2b^ko/+^; Pkhd1^del3-67/+^* trans-heterozygous mice had an intermediate phenotype (Figure 6j). As predicted, *Tfap2b* and AP-2β expression were absent from corneal endothelium and stroma (Figure 6k, l), and the AP-2β staining pattern in the eye at E13.5 was the same as that of *Pkhd1^del3-67/del3-67^* (Supplementary Figure 12e). Interestingly, none of the mice developed cystic kidney or liver disease (Supplementary Figure 12f, g), suggesting that *Pkhd1* deletion does not cause global *Tfap2b* dysregulation.

**Fig. 6:**
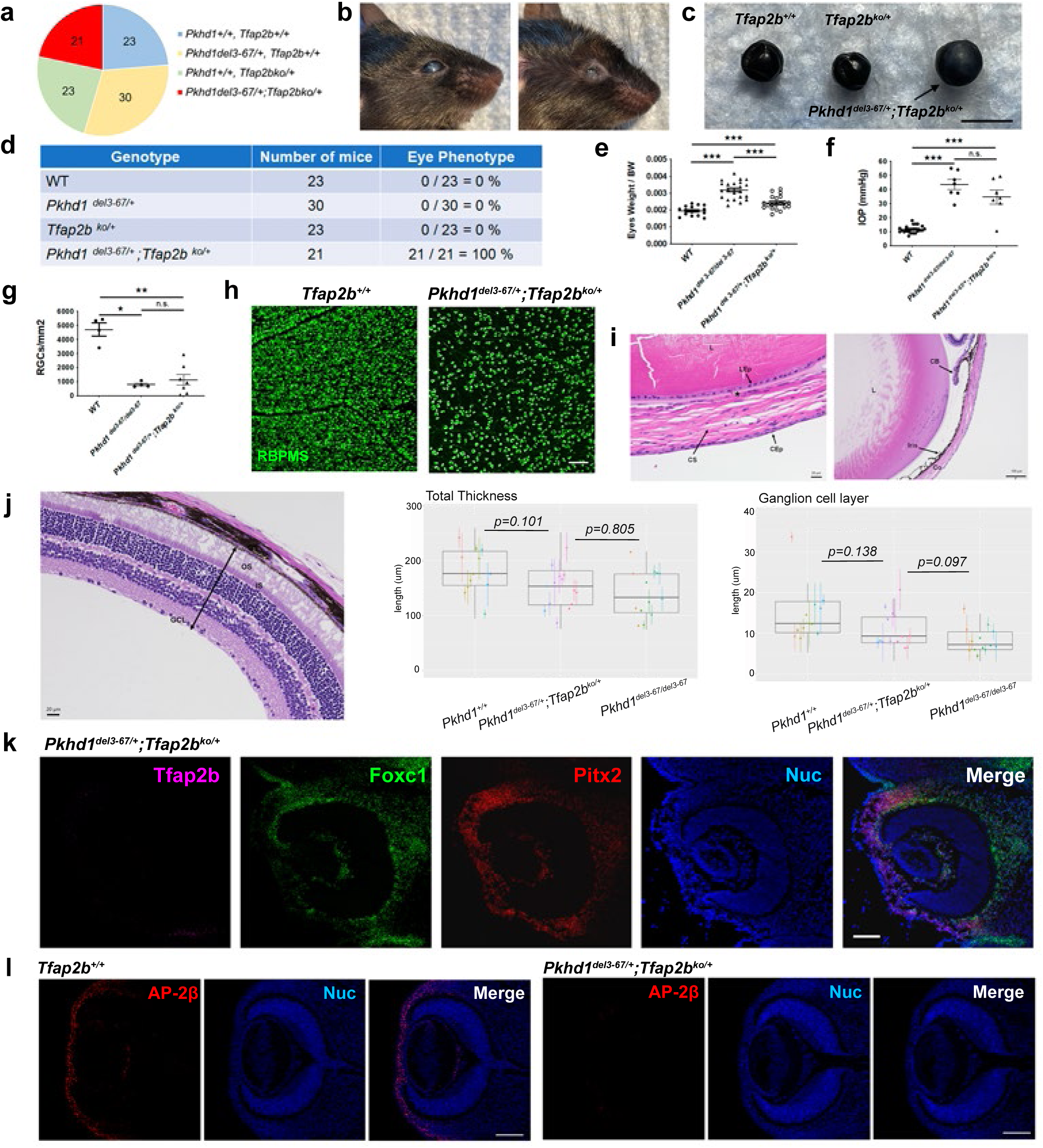
Eye phenotype of *Pkhd1^del3-67/+^;Tfap2b^ko/+^* trans-heterozygous mice. (a) Genotyping results for 97 weaned pups from 12 litters from *Pkhd1^del3-67/+^* x *Tfap2b^ko/+^* mating. (b) Representative eye images of two *Pkhd1^del3-67/+^;Tfap2b^ko/+^* trans-heterozygotes at 6 months of age. Left: representative corneal opacity, right: representative atrophic recessed eye with scarred eyelids. (c) Representative image of eye globes at 3 months of age. Scale bar 4 mm. (d) Prevalence of eye phenotype in pups from *Pkhd1^del3-67/+^* x *Tfap2b^ko/+^* mating. (e) Eyes weight per body weight (BW) at 4 weeks of age. Each dot is the mean value of both eye weights for a single mouse. ^★★★^ p < 0.001. (f) IOP of WT, *Pkhd1^del3-67/del3-67^* and trans-heterozygotes at 3 weeks of age. IOP *Pkhd1^del3-67/del3-67^* values are from the same dataset used in Figure 2a. Each dot is the mean value of IOP for a single mouse. Combined IOP data of all mouse genotypes are shown in Supplementary Figure 11c. ^★★★^ p<0.001, n.s. not significant. (g) Number of RGC of all mouse genotypes at 4 weeks of age. Each dot represents the mean number of RGC counted from a series of immunostained images from a single mouse. ^★★^ p<0.01, ^★^ p<0.05, n.s. not significant. (h) Representative images of RGC at 4 weeks of age stained with RBPMS. Scale bar 100μm. **(i) Eye pathology of** *Pkhd1^del3-67/+^*; *Tfap2b^ko/+^* trans-heterozygote with H&E staining at 1 month of age. (Scale bar 20 μm on left, 100 μm on right). CS: corneal stroma, CEp: corneal epithelium, CEn: corneal endothelium, L: Lens, CB: ciliary body, Co: cornea, LEp: lens epithelium. (j) On left, H&E-stained retina of 1 month-old *Pkhd1^del3-67/+^*; *Tfap2b^ko/+^* trans-heterozygote. Middle/Right: Box plots showing total thickness and thickness of RGC layer of retinas in *Pkhd1^del3-67/del3-67^* (n=7), *Pkhd1^del3-67/+^*; *Tfap2b^ko/+^* (n=7) and *Pkhd1^+/+^* (n=6) 1-month-old mice. Each dot is the average thickness and the range indicates maximum and minimum values per eye; the left/right eyes of an animal are identified using dots of the same color. The bars show Wilcoxon rank sum p values comparing thickness of (left) whole retina; (right) ganglion cell layer. (k) Representative images of *in situ* hybridization of Tfap2b in E13.5 eyes using RNAScope HiPlex v2. Scale bar 50 μm. (l) E13.5 mouse eye immunostained for AP-2β (red) and nuclei stained with Hoechst 33342 (blue). Scale bar 100 μm.

One surprising finding was that nearly all *Tfap2b^ko/+^; Pkhd1^del3-67/+^* trans-heterozygous mice also developed incisor malocclusion (Supplementary Figure 13a, b). We had previously reported that nearly all *Pkhd1^del3-67/del3-67^* mice develop malocclusion (Supplementary Figure 13c, d), but the sole surviving *Tfap2b^ko/ko^* mouse in our study did not have this finding (Supplementary Figure 13e). A similar phenotype had also not been reported for other *Tfap2b^ko/ko^* mice ^13,14,51,52^ or mice with *Tfap2b* deleted in cranial NCCs ^16,53,54^ However, a recent study using lineage tracing reported that *Tfap2b* is a marker for progenitors in tooth development ^55^, and a heterozygous mutation of *TFAP2B* has been associated with a variety of dental anomalies in a Thai population ^56^, suggesting a causal link between disrupted *Tfap2b* expression and incisor malocclusion observed in our study.

*In toto*, these results suggest that the *Pkhd1^del3-67^* deletion complements the mutant *Tfap2b^ko^* allele by disrupting expression of the adjacent *Tfap2b*^+^ allele in a limited set of eye-specific NCC derived cells, supporting the hypothesis that the large genomic deletion disrupts the activity of remote enhancer elements that control *Tfap2b* expression in a highly restricted manner.

## DISCUSSION

Our data suggest that the genomic architecture within or near the *Pkhd1* locus controls *Tfap2b* expression in a highly restricted set of NCC-derived cells. The large-scale deletion we generated likely disrupted TADs between *Pkhd1* and *Tfap2b* that control *Tfap2b* expression in the POM, resulting in their loss of *Tfap2b* expression (Figure 7a, b). We speculate that the same mechanism likely explains the similar phenotype previously described in mice with a complex genomic alteration induced by transgenesis ^57^, Barzago *et al* had set out to make a transgenic mouse expressing a mutant form of xanthine dehydrogenase (XDH) but inadvertently had created a new mouse model of glaucoma when their transgene random insertion caused a complicated genomic rearrangement/deletion affecting the interval between *Tfap2b* and *Pkhd1* that resulted in loss of AP-2β expression. Our model, directly targeting the *Pkhd1* locus, also nicely explains how *Pkhd1/Tfap2b* trans-heterozygous mutants develop the phenotype. The *Tfap2b^ko^* allele produces no functional transcript whereas the allele with genomic deletion of *Pkhd1^del3-67^* disrupts cis-acting genomic interactions necessary for *Tfap2b* expression from the *Tfap2b* wild type allele (Figure 7c).

**Fig. 7:**
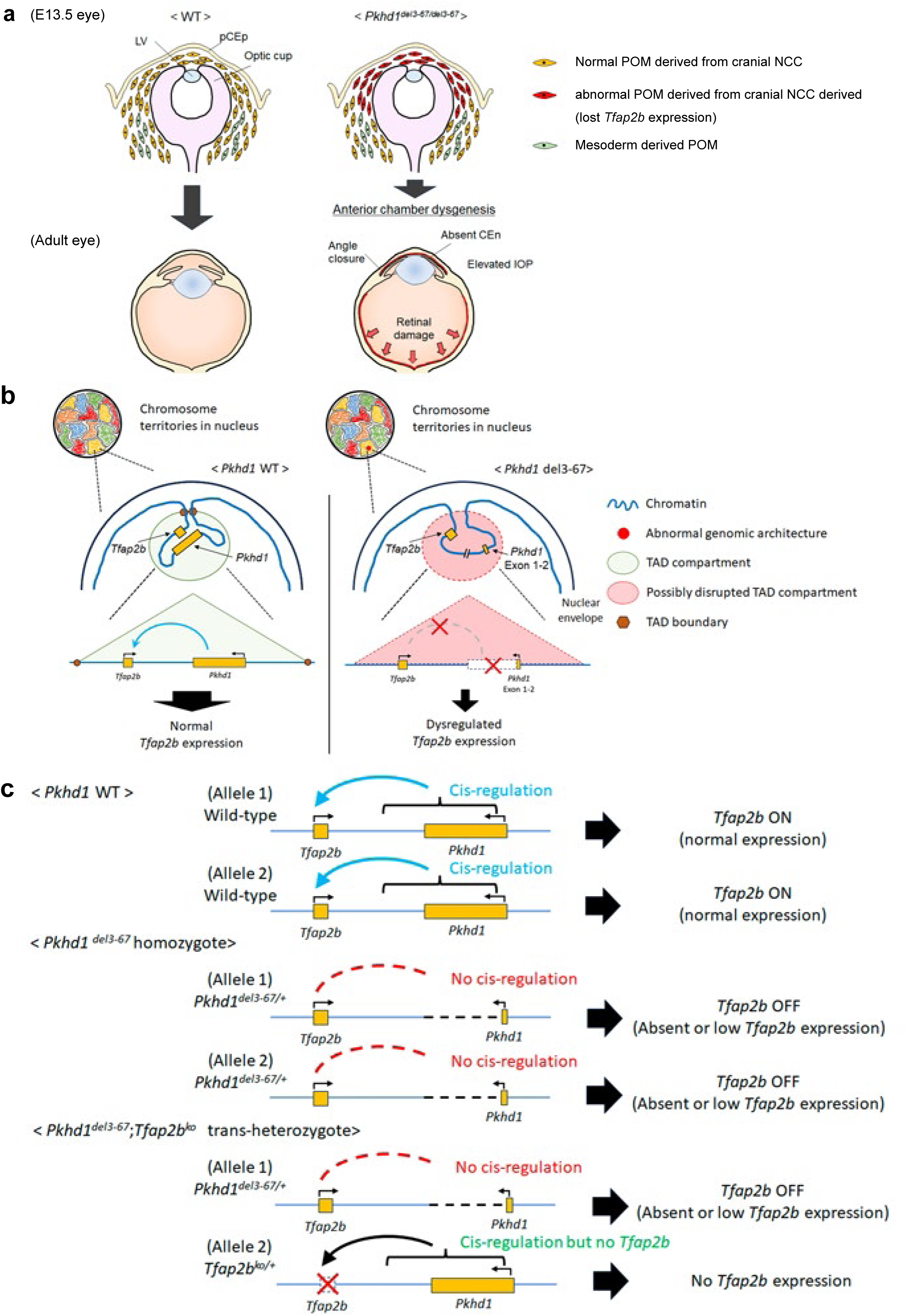
Potential model of how *Pkhd1^del3-67/del3-67^* and *Pkhd1^del3-67/+^;Tfap2b^ko/+^* trans-heterozygotes develop early onset glaucoma via *Tfap2b* down regulation. (a) In the normal E13.5 eye (left), NCC-derived POM expressing *Tfap2b* migrate to the area of the future anterior angle and anterior chamber and differentiate into corneal endothelium and stroma, the trabecular meshwork, iris stroma and ciliary body stroma. In *Pkhd1^del3-67/del3-67^* and *Pkhd1^del3-67/+^;Tfap2b^ko/+^* mutants, POM lack *Tfap2b* expression and fail to develop a normal angle or corneal endothelial layer. This results in a closed anterior angle, which over time results in increased IOP and subsequent retinal damage. (b) Cartoon showing how large deletions in the *Pkhd1* locus or nearby regions might disrupt TAD architecture and result in reduced *Tfap2b* expression in a small set of NCC-derived cells. (c) Schematic showing the likely effects of different genotypes used in this study on *Tfap2b* expression.

The initial goal of this study was to determine whether the complex pattern of splicing reported for the *Pkhd1* gene had functional significance. We did this by comparing the phenotypes of previously reported mouse lines with small deletions along the *Pkhd1* gene with a line we generated by crossing two conditional *Pkhd1* mouse lines, one with exons 3 and 4 flanked by lox-P sites and another with floxed exon 67, and expressing Cre-recombinase in the germline. This strategy resulted in a moue line with the entire genomic interval from *Pkhd1* intron 2 to the gene’s 3’UTR deleted ^27^. While the large-scale deletion did not result in any obvious differences with respect to the kidney or liver, it did yield unexpected effects on fertility ^27^ and eye development, the latter of which is described in this report.

We considered whether the phenotype was an uncommon and unrecognized feature of ARPKD, especially given that *PKHD1* encodes a ciliary protein and mutation of other ciliary genes results in eye abnormalities. In support of this hypothesis, we identified in publicly available scRNA databases a disproportionally high number of reads in a few exons, suggesting the possibility of a previously unknown *Pkhd1* transcript unique to corneal endothelial cells that could be associated with the phenotype. None of the other mutant *Pkhd1* mouse models have genomic deletions that completely remove this segment of *Pkhd1*, so this could possibly have explained their lack of a phenotype. However, an extensive literature search and personal communications with investigators who lead four large registries of individuals with ARPKD failed to identify any with relevant eye abnormalities. It is important to note that large bilineal deletions similar in size to that of the *Pkhd1^del3-67^* allele or disrupting the putative alternative *Pkhd1* transcript have not been reported in humans ^58^, so it is unknown whether humans would develop the same abnormalities if this segment of the gene was similarly deleted.

While we were unable to find evidence linking eye abnormalities to ARPKD in humans, we did find GWAS studies linking *PKHD1* to POAG. We also found another gene associated with an ARPKD phenotype in mice linked to POAG by GWAS (*Bicc1*) (Supplementary Table 2). In reviewing the relevant data for *PKHD1*, we noted that inactivation of the adjacent gene, *Tfap2b*, in NCC using a Wnt1-Cre had been previously reported to result in a phenotype like that of the *Pkhd1^del3-67/del3-67^* mutants. We also found that *Tfap2b* was highly expressed in the same adult corneal endothelial cells as *Pkhd1.* This finding was consistent with a prior study reporting that *Pkhd1* was under *Tfap2b* regulation in the kidney, suggesting a potentially relevant functional relationship between *Pkhd1* and *Tfap2b* in the eye ^36^. A similar phenotype was also reported in mice in which *Ift88*, which encodes a ciliary protein essential for normal ciliary trafficking, was conditionally inactivated in NCC cells using Wnt1-Cre ^35^. We therefore wondered if the similarities could be explained by a common feature—loss of fibrocystin function in the developing eye. This led us to hypothesize that *Tfap2b* was upstream of an eye-specific *Pkhd1* transcript, and that loss of the *Pkhd1* transcript was responsible for the mutant phenotype. Consistent with this model, potential AP-2β binding sites mapped near the putative alternative *Pkhd1* transcript. If this model were true, it would suggest that a *Pkhd1*-derived product has an essential function in eye development.

The data, however, do not support this hypothesis. We were unable to detect *Pkhd1* full-length or alternative transcripts in relevant cell types in the normal eye and we instead found tissue-specific loss of *Tfap2b* and AP-2β protein expression in neural crest cells and their derivatives in the POM region of the developing eye of *Pkhd1^del3-^*^67^*^/del3-67^* and *Pkhd1^del3-67/+^; Tfap2b^ko/+^* mutants. Importantly, we used the same AP-2β antibody for all developmental stages (E8 – E13.5), which excludes differences in antibody sensitivity or specificity as a trivial explanation for why AP-2β was detected in some but not all mutant tissues, and we confirmed that the loss of AP-2β-protein expression correlated with loss of *Tfap2b* transcript. Taken together, the data support our hypothesis that gene expression of *Tfap2b* is tightly regulated by at least two spatiotemporal tissue-specific enhancers and that cis-acting genomic interactions with at least one of them is disrupted by the *Pkhd1* del3-67 deletion.

Genome-wide association studies (GWAS) have been extremely successful in identifying thousands of loci for numerous diseases and medical conditions. For some diseases, the studies have identified loci that are linked to pathways previously implicated in the disease or condition, and demonstrating the variant’s effects on gene activity or expression is relatively straight-forward ^59^. The true “super-power” of GWAS studies, however, is in their ability to identify novel or unsuspected factors that either contribute to disease or protect against it, but the challenge can be in establishing how the variant does so. This is particularly true for single nucleotide polymorphisms that lie outside of known genes or regulatory regions.

Primary open angle glaucoma is a relevant example. Iterative GWAS studies have identified an ever-growing list of loci linked to the condition or its endophenotypes. While several loci are in genes with clear links to the pathobiology of the disease, the vast majority are not, including those close to the *PKHD1* locus. In fact, none of the SNPs linked to POAG in Figure 1b were reported as linked to *TFAP2B* in a recent publication that describes over 200K long range GWAS SNP-target gene interactions ^60^. In this study, we provide *in vivo* evidence showing that genomic elements within or near *Pkhd1* control *Tfap2b* expression in a very limited set of neural crest-derived cells in the POM, and the latter is responsible for the corneal endothelial and stromal abnormalities that were observed. There is precedence for similar mechanisms in humans: a recent report describes a patient with branchio-oculo-facial syndrome caused by NCC developmental defects due to an inversion that separated *TFAP2A* from its enhancer ^61^.

We posit that a similar process likely explains why SNPs near the *PKHD1* locus are linked to glaucoma in humans. In mice, the large *Pkhd1* exon 3-67 deletion disrupts TADs that link the loci whereas in humans the SNPs likely have more subtle effects. Many of the SNPs identify protective alleles, which perhaps exert their effects by enhancing or maintaining *TFAP2B* expression in target cells. Interestingly, a recent study reported that SNPs linked to distant genes had significantly more insertions and/or deletions around them than other SNPs ^62^. This suggests the possibility that human *PKHD1* SNPs linked to POAG are markers for nearby genomic alterations that affect TADs which regulate *TFAP2B* activity.

At first glance, it might seem surprising that in mice altering *Tfap2b* expression results in anterior segment dysgenesis with a closed irido-corneal angle and early onset glaucoma while in humans the disease is late onset and open angle. However, several other genes associated with POAG in humans listed in Figure 1a also result in congenital glaucoma with angle closure ^63^ and abnormal retinal morphology (www.mousephenotype.org) ^64^ when deleted in mice, suggesting this is a common pattern. In the case of *ANGPT1*, heterozygous inactivating mutations in humans result in primary congenital glaucoma while SNPs are associated with POAG ^63^. It is also worth noting that keratoconus, a defect of the cornea that is a common indication for corneal transplantation, also is associated with SNPs in the *TFAP2B/PKHD1* interval. They are distinct from those associated with POAG and increase the risk of disease ^65^. Collectively, these findings suggest that several pathways may be common to the different conditions, with the severity and timing of disease determined by when and how much the pathways’ functions are dysregulated.

It also might be surprising that the corneal epithelium of our *Pkhd1^del3-67/del3-67^* and *Pkhd1/Tfap2b* trans-heterozygous mutants was abnormal given that it is derived from the surface ectoderm and not the POM. A similar finding was reported for mice with *Tfap2b* conditionally inactivated in NCCs ^16^. The investigators subsequently showed that corneal epithelial cell fate was perturbed in these mutants and hypothesized that it was due to altered regulatory signaling from the POM-derived mutant stroma. Prior studies also have shown that loss of the corneal endothelium can result in reduced corneal stratification ^66,67^ which is consistent with their hypothesis. Altered stromal-epithelial cell signaling may also explain the pathology in keratoconus and how variants in the *PKHD1/TFAP2B* are linked to both POAG and keratoconus ^68^.

Craniofacial abnormalities are frequently associated with *TFAP2B* mutations in mice and humans, but isolated dental anomalies are uncommon. While incisor malocclusion was not reported for any other mouse model with *Tfap2b* inactivation, it is possible that differences in genetic background, under reporting, analyses restricted to young mice, or an effect specific to the *Pkhd1^del3-^*^67^ allele account for the difference. However, given that *Tfap2b* precursors give rise to teeth, the mechanism of their malformation in our mouse models almost certainly results from the disruption of *Tfap2b* activity in a restricted set of cells.

In conclusion, while our search for a relationship between ARPKD and eye phenotypes in humans was unsuccessful, we did find strong evidence for a functional relationship between the *PKHD1* locus and eye disease in humans. Our studies suggest that the SNPs near *PKHD1* associated with POAG in GWAS studies likely cause disease by altering the transcriptional activity of *TFAP2B* in a highly defined set of neural-crest derived cells. Our studies further hint at the complex developmental patterns of regulatory control of *Tfap2b*.

## METHODS

### Mice

The *Pkhd1^del3-67^* mouse line was described previously ^27^. The *Tfap2b* knockout mouse line was previously described ^16,52^ and kindly provided by Drs Trevor Williams (University of Colorado) and Judith A West-Mays (McMaster University Health Science Centre). Littermates were used for each experiment and were generally the offspring of *Pkhd1 ^del3-67/+^* x *Pkhd1 ^del3-67/+^* or *Pkhd1 ^del3-67/+^* x *Tfap2b^ko/+^* mating, though we also used *Pkhd1 ^del3-67del3-67+^* (female) x *Tfap2b^ko/+^* (male) mating to increase the probability of getting trans-heterozygotes. Because the background of the *Pkhd1^del3-67^* mouse line is C57BL/6N, we excluded the *rd8* mutation (a single nucleotide deletion in *Crb1*), which is common in that strain, in our mice ^69^. For eye weights, we compared the mean values for all eyes of each genotype, but we used the mean values for both eyes as a single value in the eyes weight/body weight analysis. Body weight (BW), eye weight and BW/Eye weight data are presented as the means ± standard error. Data for the two groups were analyzed by two-sided Mann-Whitney test using Prism 10 software. The *Pkhd1^del3-^*^67^ mouse line is available upon request.

### Genotyping

Genomic DNA was isolated and genotyped using the REDExtract-N-Amp tissue PCR Kit (SIGMA) according to the manufacturer’s protocols. *Pkhd1^+^*, *Pkhd1^del3–67^* and *Tfap2b^ko^* allelic products were run on 2% agarose gels. Primer information is described in Supplementary Table 1. Conditions for genotyping the *Tfap2b* knockout allele were kindly provided by Dr. Trevor J. Williams at University of Colorado.

### Intraocular pressure (IOP) measurement

IOP was measured using a rebound tonometer (Tonolab, Colonial Medical Supply). IOP was recorded during the same 1-hour window (9:00-10:00 AM) each day, sampled 6 times for each eye and automatically averaged for each individual recording after the elimination of the highest and lowest values. Means IOP of both right and left eye were used as IOP of the mouse. All data are presented as the means ± standard error. Data for the two groups were analyzed by two-sided Mann-Whitney test using Prism 10 software.

### Histology

Adult mouse eyes were dissected and fixed in 10% neutral buffered formalin at 4°C overnight, washed in 70% ethanol and then stored at room temperature in 70% ethanol. Whole embryonic eyes were fixed in 10% neutral buffered formalin at 4°C overnight, dehydrated in a graded alcohol series and embedded in paraffin. The pupillary-optic nerve plane was cut with vertical sections in adult eyes and coronal sections in embryos, and all sections were stained with hematoxylin and eosin (H&E).

### Immunostaining

E8 and E9 mouse embryos prepared from *Pkhd1^del3-67/+^* x *Pkhd1^del3-67/+^* mating were prepared as follows: fixed in 4% PFA for one hour at room temperature (RT); washed three times with room temperature phosphate-buffered saline (PBS); permeabilized with 0.5% PBST (PBS with 0.1% Triton x100 solution) for 20 min at RT; rinsed three times with PBS; incubated with primary antibodies at 4° C overnight; incubated for 2 h at room temperature with secondary antibodies diluted at 1:1000 and Hoechst at 1:2000; rinsed three times with PBS; and mounted on microscope slides using imaging spacers 9 mm diameter x 0.12 mm depth from grace Bio-Labs. Samples were imaged with a Zeiss 800 confocal microscope. For crossed sections, previously stained whole mouse embryos were incubated in 30% sucrose in 0.1M phosphate buffer until the embryos sank and embedded in optimum cutting temperature (OCT) medium, sectioned (10μm) on a CryoStar Cryostat (Thermo Fisher Scientific), washed 2 times with PBST (PBS-0.1% Tween20) for 5 minutes, and then mounted using Fluoromount-G (SouthernBiotech Laboratories).

E13.5 and E15.5 mouse embryos were fixed in 4% PFA for 24 hour at RT, embedded in paraffin, treated with universal HIER antigen retrieval reagent (ab108572, Abcam) according to the user guide, incubated with primary antibodies at 4℃ overnight, incubated with secondary antibodies for 1 hour at RT with Hoechst 33342 (x1000), and mounted using Prolong Glass Antifade Mountant (P36984, Thermo Fisher Scientific) Images were captured with a Nikon CSU-W1 SoRa Spinning Disk Confocal Microscope.

For studies of the retina and angle of the adult eye, dissected aphakic eyes were fixed in PFA, equilibrated in 30% sucrose in PBS overnight and frozen in O.C.T. compound (#4585, Fisher Healthcare). Frozen eyes were cut into 10 mm-thick sections using a cryostat (CM1860, Leica Biosystems). Sections that included the optic nerve head were collected on glass slides. incubated in blocking buffer (1xTBS, 0.05% Tween 20, 1% BSA, 1% horse serum and 0.02% sodium azide) for 1 hr, incubated in blocking buffer with primary antibodies and Rhodamine phalloidin (R415, Thermo Fisher Scientific) at 4°C overnight, washed for 20 mins three times with buffer (1xTBS with 0.05% tween20), incubated in secondary staining solution and DAPI in blocking buffer without horse serum at room temperature for 1 hr, washed for 20 mins with buffer and then imaged with the Zeiss 800 confocal microscope.

All images being compared in a group were processed in like manner: deconvolution or denoising, then contrast-stretching using Fiji software ^70^.

### Retinal ganglion cell (RGC) quantitation

Mice were CO_2_ euthanized and eyes were enucleated and fixed in 4% paraformaldehyde in PBS (PFA) for 10 min following the creation of a hole through the cornea with a 25G needle. The lens and the cornea were removed, four incisions were made to open the retina into a flat mount, and the retina was separated from the sclera. After fixing with PFA for an additional 20 min, washing with PBS three times for 10 min and blocking in blocking buffer containing 1xTBS with 1% Triton X-100, 1% bovine serum albumin (BSA), 1% horse serum and 0.02% sodium azide for 1hr, retinas were incubated with primary antibodies in the blocking buffer at 4°C for three days. Retinas were washed with the washing buffer (1xTBS with 1% Triton X-100) for 20 mins three times and transferred into blocking buffer with secondary antibodies and DAPI. After incubation at 4°C overnight, retinas were washed with washing buffer for 20 min three times and placed on a glass slide with mounting medium (Fluoro-Gel, Electron Microscopy Sciences). For RGC counting (RGCs/mm^2^) of whole-mount retinae, four separated fields positioned in the middle part of each retina were selected and the images were acquired by a confocal laser-scanning microscope (LSM 700, Carl Zeiss Inc.). Investigators were blinded to the identity of the samples at the time of image analysis. The number of RBPMS-positive RGCs in a rectangular area (20702 mm^2^) were counted in each image using Fiji software (National Institutes of Health) and the average RGC density (RGCs/mm^2^) was calculated. One retina was analyzed of each mouse. All data are presented as the means ± standard error of them. Data for two groups were analyzed by two-sided Mann-Whitney test using Prism 10 software.

### Cornea thickness measurements

One eye of each of three *Pkhd1^del3-67/del3-67^* and four control 3-week-old mice was embedded in paraffin, sectioned, H&E stained and imaged at x20 magnification using Keyence BZ-810. Using the line and measurement tools in ImageJ 1.54p, the width of three regions of the central cornea was measured in each animal and the animal width averages were compared using two-sided Welch t-test in R 4.2.3.

### Retinal layer measurements

Images of both right and left eyes of H&E-stained retinas for each mouse, avoiding the region surrounding the optic nerve. The images were analyzed in ImageJ ^71^, measuring the total thickness of the retina and of each of the following layers: ganglion cells, inner plexiform, inner nuclear, outer plexiform, outer nuclear, layer of rods and cones. For each layer, the average length of all measurements of each mouse were analyzed in R using Anova for multiple group comparisons and two-sided Wilcoxon rank test for two groups. Representative images were chosen from mice with total retinal thickness close to the group mean thickness.

### Antibodies

We used the following primary antibodies for immunostaining frozen specimens: anti-Sox10 (1:100, 2ug/ml) sc365692 Santa Cruz Biotechnology; anti-AP-2β (1:100) cst2509 Cell Signaling Technology; anti-Sox9 ab5535 (1:500, 0.002ug/ml, Millipore Sigma); anti-ZO-1 (1:100, 2.5ug/ml) #40-2200 Invitrogen); anti-smooth muscle actin (AB5694, 1:300, Abcam); anti-RBPMS (ABN1376, ABN1362 1:300, Millipore Sigma); anti-Pax6 (PRB-278P, 1:300, Covance). The following secondary antibodies used for frozen sections were obtained from Thermo Fisher Scientific: donkey anti-rabbit Alexa fluor 488 (1:1000), Alexa555-anti-mouse IgG, donkey anti-mouse Alexa fluor 568 (1:1000), donkey anti-rabbit Alexa fluor 647 (1:1000). Either DAPI or Hoechst 33342 were used for nuclear staining.

### Transcriptomic analysis for human and mouse eyes

Human eye transcriptomes at the single-cell level were queried to characterize expression of *PKHD1* in different cell types using the Single Cell Portal of “Cell atlas of the human ocular anterior segment: Tissue-specific and shared cell types” (Single Cell Portal (broadinstitute.org) ^42^ . For the mouse eye transcriptome, GSE183690 ^43^ and GSE158454 ^44^ datasets at the single-cell level were used to characterize expression of genes in the *Pkhd1* genomic region in different cell types. Sequence files were downloaded using sratoolkit v.3.0.5 ^72^ and processed with celllranger v.7.1.0 using mm10 mouse reference genome. Reads in the chromosome 1 genomic region chr1:20,055,779-20,620,057, which includes *Pkhd1* and non-coding RNA *4930486I03Rik* and *Gm24162*, were extracted using samtools v.1.17 ^73^. Count matrices were imported into R v.4.3.0 ^74^ and analyzed with Seurat v.4.3.0 ^75^. Cell types were assigned using cluster markers reported in ^42^ and ^43^. Reads in the BAM files that were confidently assigned to single cell tags were labeled with cell type identities in R and the results were plotted using ggplot2 v.3.4.2 ^76^. Enhancers in the *Pkhd1* genomic region were identified in UCSC Genome Browser ^77^ using mouse genome mm10 and scanned for AP2-β motifs with TFmotifView ^78^. Evidence for association between the human *PKHD1* locus and ocular phenotypes was investigated using the Common Metabolic Diseases Knowledge Portal ^41^ and supplemented with a literature review. Visualization of genomic regions was done in UCSC Genome Browser, Ensembl ^79^ or Integrative Genomics Viewer (version 2.11.9) ^80^.

### Analysis for topologically associated domains in human and mouse genome

Visualization of topologically associated domains (TADs) described in ^81,82^ was done using 3D Genome Browser ^83^. Chromatin loops inferred from experiments using mouse ^84–86^ and human ^81,87–94^ samples were downloaded from 3D Genome Browser and visualized in R with the circlize package ^95^.

### *In situ* hybridization with RNAscope HiPlex assay v2

*In situ* hybridization to detect *Pkhd1* exons was performed according to the user manual. Briefly, formalin-fixed paraffin-embedded 2 months old WT mouse eyes (fixed 24h at RT) were used as samples (N=3). Pretreatment was performed with 15 mins of protease Ⅲ at 40℃. To reduce autofluorescence, 5% RNAscope HiPlex FFPE reagent was used for each sample. An exon 54 and 55 *Pkhd1* transcript (detected in mouse single cell transcriptome data) was too short to make a specific probe for the HiPlex v2 assay; therefore, we designed a probe targeting *Pkhd1* exons 54-56 (Cat. #1281151-T3). Cat. #1281161-T6 probe was used to detect whole *Pkhd1* transcripts and a RNAscope HiPlex12 Negative Control Probe was used as a negative control. Nuclei were stained with RNAscope DAPI. Prolong Gold Antifade Mountant (P36934, Thermo Fisher Scientific) was used as mounting media. Mouse liver tissue was used as a positive control sample for *Pkhd1* detection (N=1). Images were taken using a Nikon CSU-W1 SoRa Spinning Disk Confocal Microscope with objective lens Plan Apo λ 20x/0.75. To increase the visibility of the *Pkhd1* signal we displayed the signal from the far-red channel as green in this figure. All images being compared in a group were processed in like manner: deconvolution or denoising, then contrast-stretching using Fiji software ^70^.

*In situ* hybridization to detect *Tfap2b*, *Pitx2* and *Foxc1* was also performed according to the manufacturer’s instructions. Briefly, E13.5 mouse heads were fresh frozen in OCT compound (Fisher Healthcare) and horizontally cut to 10μm thickness by a cryostat (N=1 for each genotype). The sections were collected on RNase-free glass slides and kept at -80 until use. Pretreatment was performed with 15 mins of protease at RT. Custom probes targeting mouse *Tfap2b* (Cat. #536371-T3) and POM markers genes (*Pitx2*: Cat. #412841-T2 and *Foxc1*: Cat. #412851-T1) were designed and produced by the manufacturer. Nuclei were counterstained with DAPI, and sections were mounted with ProLong Gold Antifade Mountant (P36934, Thermo Fisher Scientific). Images were acquired using a Zeiss, LSM-880 microscope.

### Whole Genome Sequencing

As described in our original report of the *Pkhd1^del3-^*^67^ line ^27^, we generated 4 independent *Pkhd1^del3-^*^67^ founders and bred them separately to homozygosis. All independently generated lines had overlapping phenotypes, including the eye anomalies, and were subsequently intercrossed. To further confirm that the line did not carry additional changes other than the intended deletion, we performed whole genome sequencing (WGS) of a homozygous *Pkhd1^del3-^*^67*/del3-*67^ mouse using Illumina Platform, 150bp paired-end. Reads were processed using BWA-MEM2 ^96^ and samtools ^97^. Visual inspection using Integrative Genomics Viewer ^80^ confirms absence of reads in the intended genomic region (chr1:20,053,820-20,613,713) and presence of reads in nearby genomic intervals, including the *Tfap2b* genomic region (Supplementary Figure 9). Structural variation analysis using CNVnator ^98^ identifies a large genomic deletion in the *Pkhd1* locus but no changes in the *Tfap2b* locus or in the region between these two loci.

### Sex as a biological variable

Our study examined male and female animals and found no differences at the time points studied. The results are described in the text and presented in the supplemental figures. We have also included the sex of the animal for specimens in the data files that will be publicly shared with the report.

### Study Approval

Mouse studies were performed under protocols K001-KDB-19, K001-KDB-22, and NEI-556 which had been approved by the NIH Animal Care and Use Committee. Mice were housed in pathogen-free animal facilities accredited by the American Association for the Accreditation of Laboratory Animal Care and meet Federal (NIH) guidelines for the humane and appropriate care of laboratory animals.

## DATA AVAILABILITY

All data and relevant metadata described in the results or presented in the figures and supplementary figures of this study have been deposited in Figshare with the following accession numbers 10.6084/m9.figshare.25970203 and will be available immediately once the report is published.

## AUTHOR CONTRIBUTIONS

YI designed research studies, conducted experiments, acquired data, analyzed data and co-wrote the manuscript. LFM designed research studies, analyzed data and co-wrote the manuscript. NN, BSK, YH, TY, JR, and FZ helped conduct experiments, acquire and analyze data. ST and LK helped design research studies. GGG designed research studies, reviewed the data and co-wrote the manuscript.

## ACKNOWLEDGEMENTS

This work utilized the computational resources of the NIH HPC Biowulf cluster. (http://hpc.nih.gov) and the NIDDK Advanced Light Microscopy and Image Analysis Core. We would like to thank Dr. Trevor Williams (University of Colorado) and Dr. Judith A West-Mays for the providing the *Tfap2b^ko^*mouse line and for helpful discussions; Dr. Lisa Guay-Woodford (Children’s Hospital of Philadelphia), Dr. Max Liebau (University Hospital of Cologne), Dr. Arlene Chapman (University of Chicago School of Medicine) and Dr. Neera Dahl (Mayo Clinic) for searching for eye phenotypes in their child and adult ARPKD patient cohorts; and Dr Elissa Lei (National Institutes of Diabetes and Digestive and Kidney Disease) for discussing potential 3-dimentional organization of DNA interaction between *Pkhd1* and *Tfap2b*. This work was supported by the intramural program of the National Institute of Diabetes and Digestive and Kidney Diseases ZIA DK075042, National Eye Institute (ZIA EY000311) and the National Institute of Dental and Craniofacial Research (ZIA DE000748)

---

**Supplementary Fig. 1:**
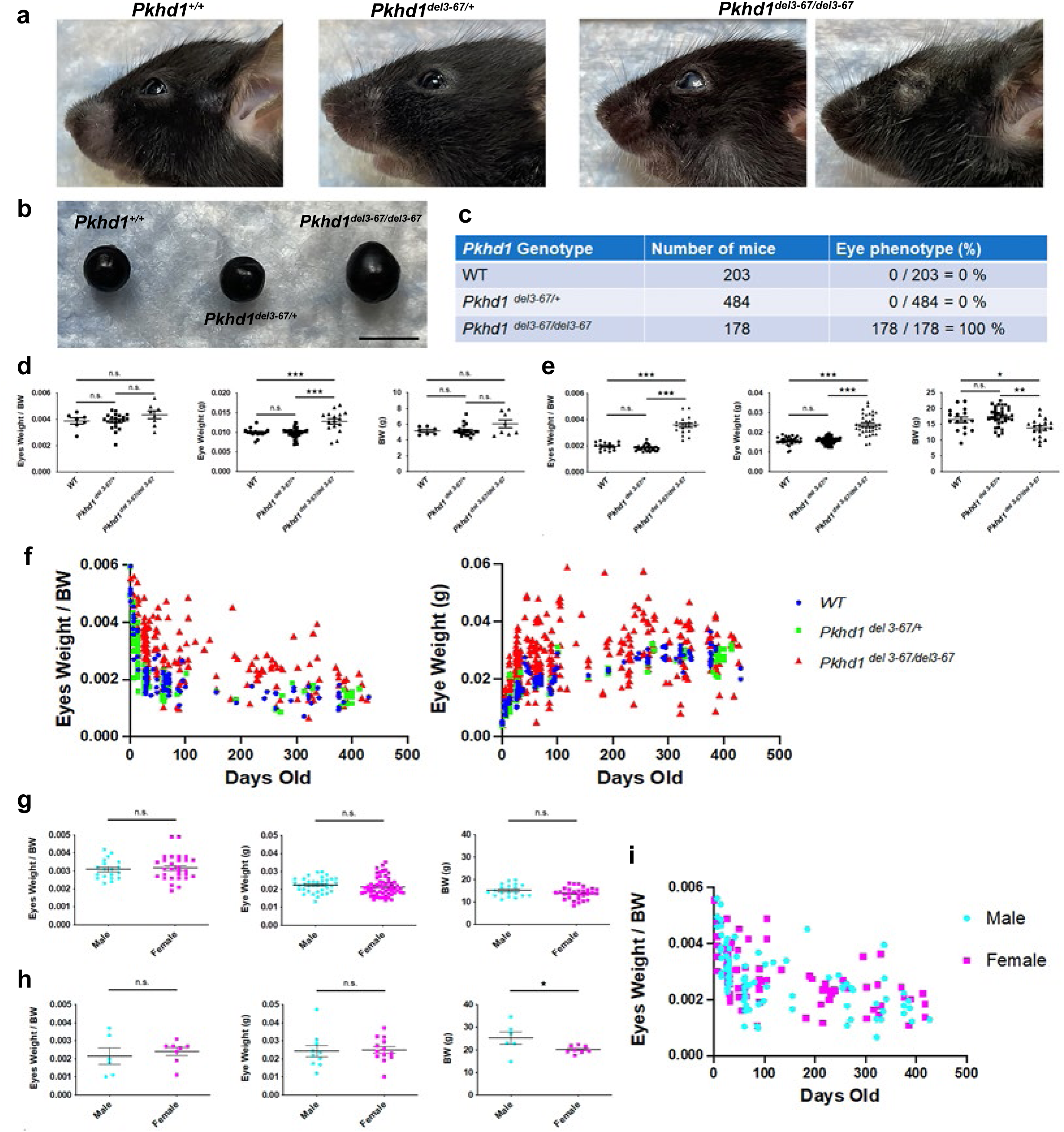
Characterization of the eye phenotype of *Pkhd1^del3-67/del3-67^* mutants. (a) Representative images of eyes from *Pkhd1^del3-67/del3-67^* and littermate controls at 2 months of age. All *Pkhd1^del3-67/del3-67^* mice developed corneal opacity but as shown in the right some of the mice develop atrophic recessed eyes with scarred eyelids as they age. (b) Representative eye globes of *Pkhd1^del3-67^* mouse line at 3 months of age. All mice are littermates. (c) Prevalence of eye phenotype in the *Pkhd1^del3-67^* mouse line. (d) From left, eyes weight per body weight, eye weight and BW at 1 week of age. For Supplementary Figure 1d-i, each dot in Eyes Weight/BW is the mean value of eyes from a single mouse while each dot in eye weight indicates the measurement for a single eye. in Eye Weight indicates a single eye. ^★★★^ p < 0.001, ^★★^ p < 0.01, ^★^ p<0.05. n.s. not significant). (f) Eye size at different ages of *Pkhd1^del3-67^* mouse line. left; scatter plots of eyes weight per BW and right; eye weight at different ages. (g) Comparison of eye size of *Pkhd1^del3-67del3-67^* male and female mice at around 1 month of age. From left: eyes weight per BW, eye weight and BW. There are no significant differences in these parameters. (h) Comparison of eye size of *Pkhd1^del3-67del3-67^* male and female mice at around 3 months of age. From left: eyes weight per BW, eye weight and BW. Mice are 90 ± 3 days old mice. ^★^ P < 0.05. n.s. not significant. (i) Scatter plot of eyes per BW of Pkhd1del3-67del3-67 of male and female mice. There is no evidence to suggest there is a significant difference in eye phenotype of *Pkhd1^del3-67del3-67^* based on sex.

**Supplementary Fig. 2:**
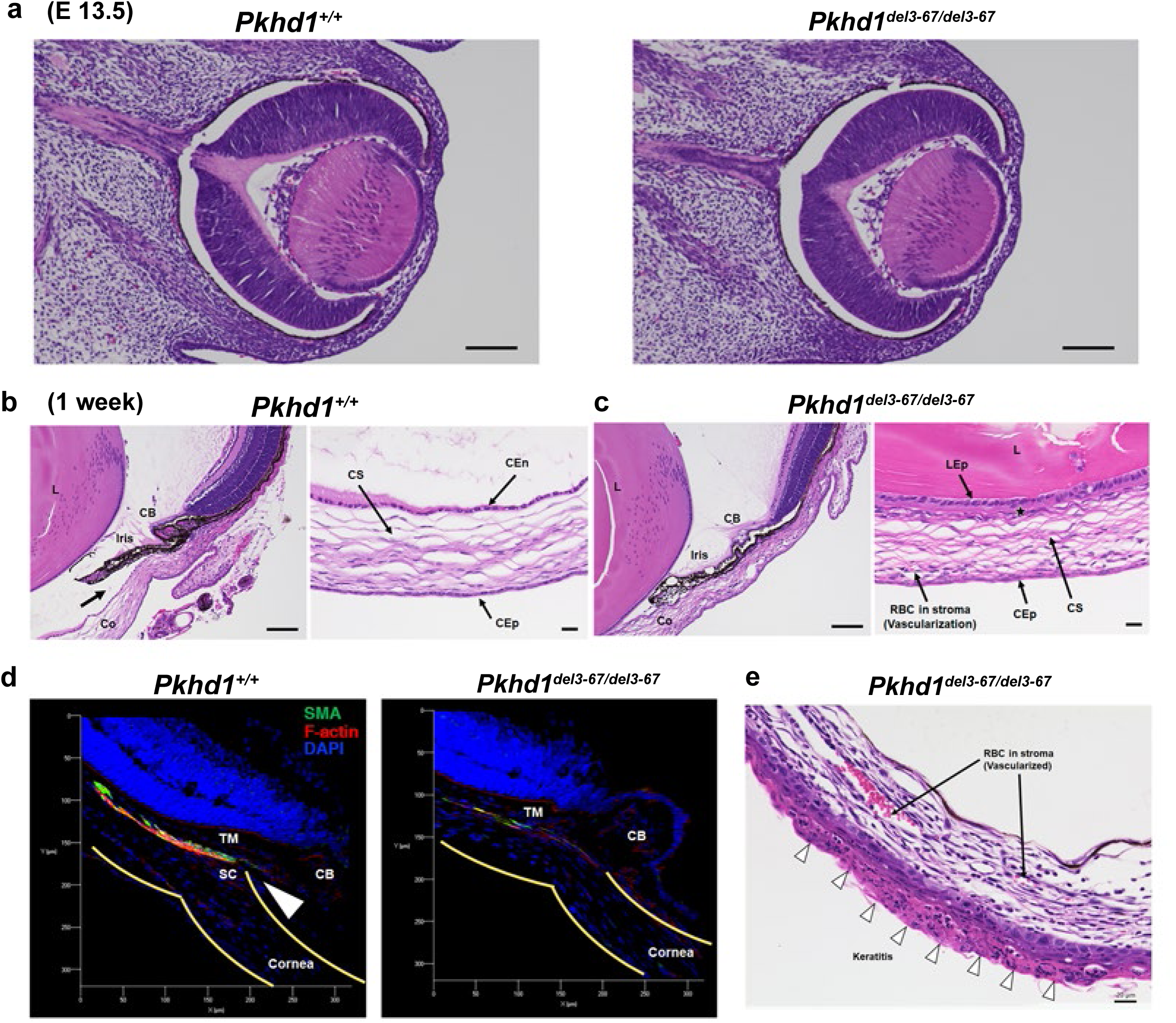
Additional data for *Pkhd1^del3-67/del3-67^* mouse eye pathology. (a) Representative H&E staining pattern of E13.5 control (left) and littermate (right) *Pkhd1^del3-67/del3-67^* eyes showing no differences. Scale bar 100 μm. (b, c) Representative eye pathology at 1 week of age (H&E staining). Left: angle tissue with 100 μm scale bar; Right: cornea with 20 μm scale bar. (b) Littermate control. The arrow in the left panel identifies the open angle in the control which is absent in the mutant. (c) *Pkhd1^del3-67/del3-67^* mutants also have abnormal ciliary body, lack a corneal endothelium, and have a vascularized corneal stroma. CB: ciliary body, Co: cornea, CEn: corneal endothelium, CS: corneal stroma, CEp: corneal epithelium, L: lens, LEp: lens epithelium. The star indicates absent corneal endothelium in *Pkhd1^del3-67/del3-67^*. (d) Immunofluorescence of the angle of the anterior chamber at 4 weeks of age. Left: control, right: *Pkhd1^del3-67/del3-67^*. The arrowhead indicates the angle of the anterior chamber, which is closed in *Pkhd1^del3-67/del3-67^*. TM: trabecular meshwork, SC: Schlemm’s canal, CB: ciliary body. Yellow line indicates surface of the eye and cornea. Scale is shown in μm. (f) 1 month old *Pkhd1^del3-67/del3-67^* eye with keratitis and obvious vascularization. Arrow heads indicate keratitis.

**Supplementary Fig. 3:**
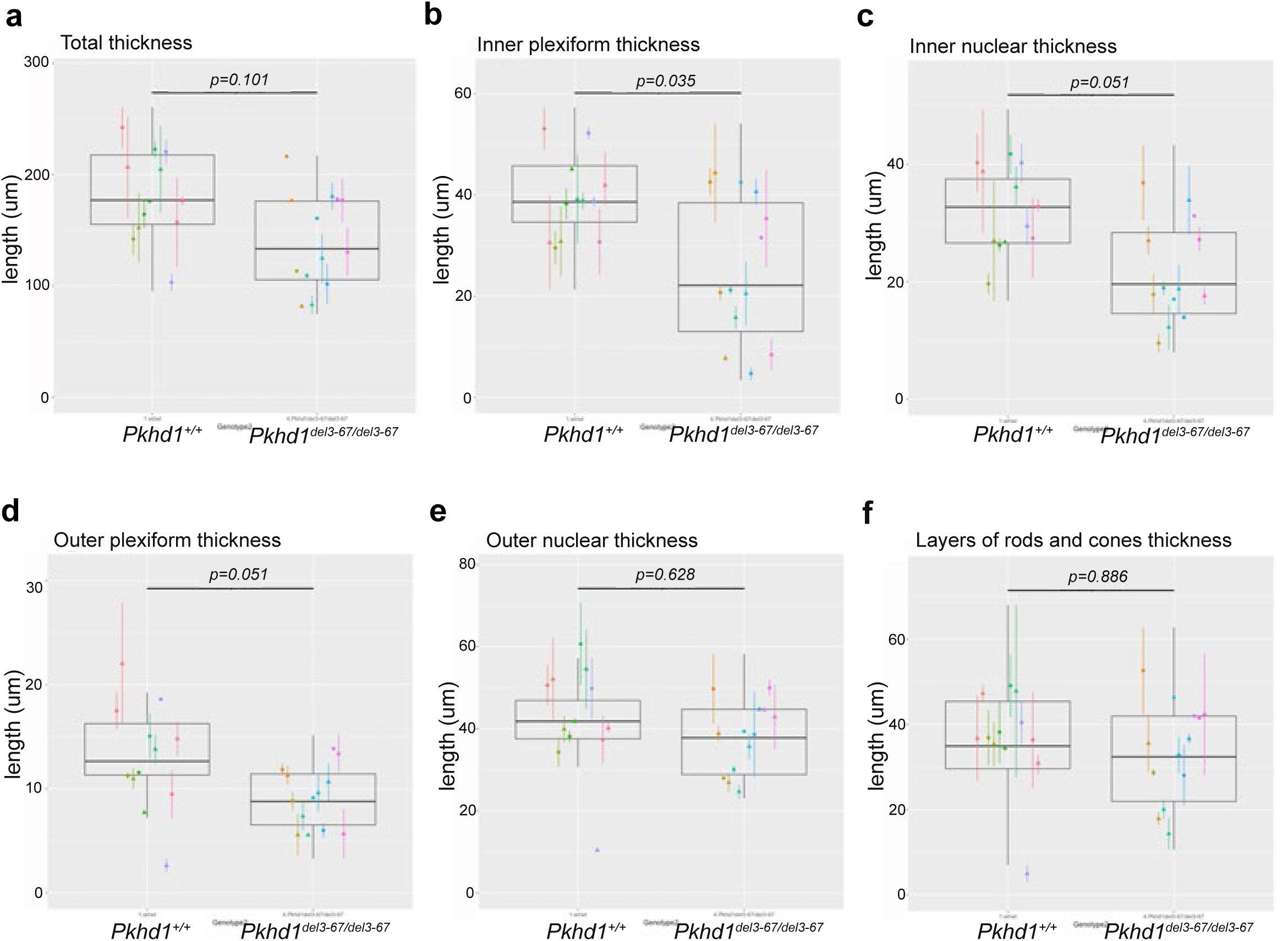
Evaluation of retinal layers of *Pkhd1^del3-67/del3-67^*. Box plot showing thickness of layers of retina in *Pkhd1^del3-67/del3-67^* (n=7) and *Pkhd1^+/+^* (n=6) 1-month-old mice. (a) Total retina, (b) inner plexiform layer, (c) inner nuclear layer, (d) outer plexiform layer, (e) outer nuclear layer, (f) rods and cones. Each dot is the average thickness and the range indicates maximum and minimum values per eye; the left and right eye of a single mouse are colored the same. The bars show Wilcoxon rank sum p-values.

**Supplementary Fig. 4:**
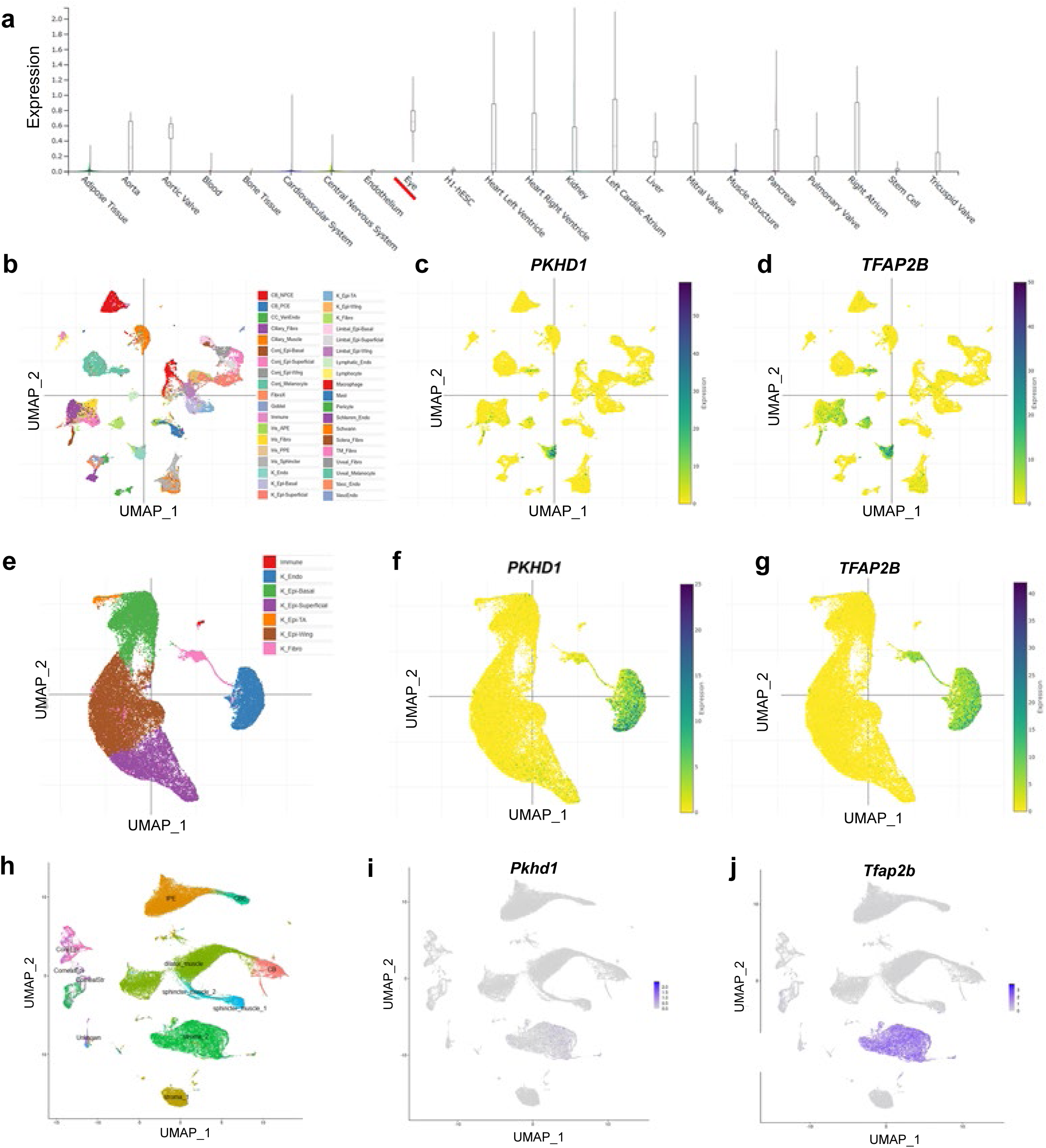
Transcriptomic analysis of human and mouse eye. (a) Tissue-specific *PKHD1* transcripts expression from the Common Metabolic Diseases Knowledge Portal. Median *PKHD1* expression level is highest in the eye. (b-d) UMAP plots of adult human eye data of integrated cell type clusters. (b) clusters included all cell types; CB_NPCE: ciliary body nonpigmented ciliary epithelium, CB_PCE: ciliary body pigmented ciliary epithelium, CC_VenEndo: collector channel/venous endothelium, Ciliary Fibro: ciliary fibroblast, Conj_Epi_Basal: conjunctival epithelium basal, Conj_Epi_Superficial: conjunctival epithelium superficial, Conj_Epi_Wing: conjunctival epithelium wing, Conj_Melanocyte: conjunctival melanocyte, FibroX: transcriptomically similar to scleral fibroblast but exhibit distinct markers, Goblet: goblet cell, Immune: immune cell, Iris_APE: iris anterior pigmented epithelium, Iris_Fibro: iris fibroblast, Iris PPE iris posterior pigmented epithelium:, K_Endo: corneal endothelium, K_Epi_Basal: corneal epithelium basal, K_Epi_Superficial: corneal epithelium superficial, K_Epi_TA: corneal epithelium transit amplifying, K_Epi_Wing: corneal epithelium wing, K_Fibro: corneal fibroblast, Limbal_Epi_Basal: limbal epithelium basal, Limbal_Epi_Superficial: limbal epithelium superficial, Limbal_Epi_Wing: limbal epithelium wing, Lymphatic_Endo: lympatic endothelial, Mast: mast cell, Schlemm_Endo: Schlemm canal endothelium, Schwann: Schwann cell, Sclera_Fibro: sclera fibroblast, TM_Fibro: trabecular meshwork fibroblast, Uveal_Fibro: uveal fibroblast, Uveal_Melanocyte: uveal melanocyte, Vasc_Endo: vascular endothelium, VascEndo: transcriptomically similar to vascular endothelium but exhibit distinct markers. (c) *PKHD1* expression pattern in all clusters. *PKHD1* expression is mostly in the “corneal endothelial cell” cluster (dark: higher expression); (d) *TFAP2B* expression in all clusters. *TFAP2B* expression is highest in corneal endothelial cells, fibroblasts, and conjunctival melanocytes. (e-g) UMAP plots of adult human eye data restricted to cell types present in the corneal sub-cluster. This figure is derived from the same dataset used in Supplementary Figure 4b-d, which included all cell types. Cell types present in the corneal sub-cluster: Immune: immune cell, K_Endo: corneal endothelium, K_Epi_Basal: corneal epithelium basal, K_Epi_Superficial: corneal epithelium superficial, K_Epi_TA: corneal epithelium transit amplifying, K_Epi_Wing: corneal epithelium wing, K_Fibro: corneal fibroblast. (h-j) UMAP plots of adult mouse eye data showing (h) cell type clusters, *Pkhd1* (i), and *Tfap2b* expression (j).

**Supplementary Fig. 5:**
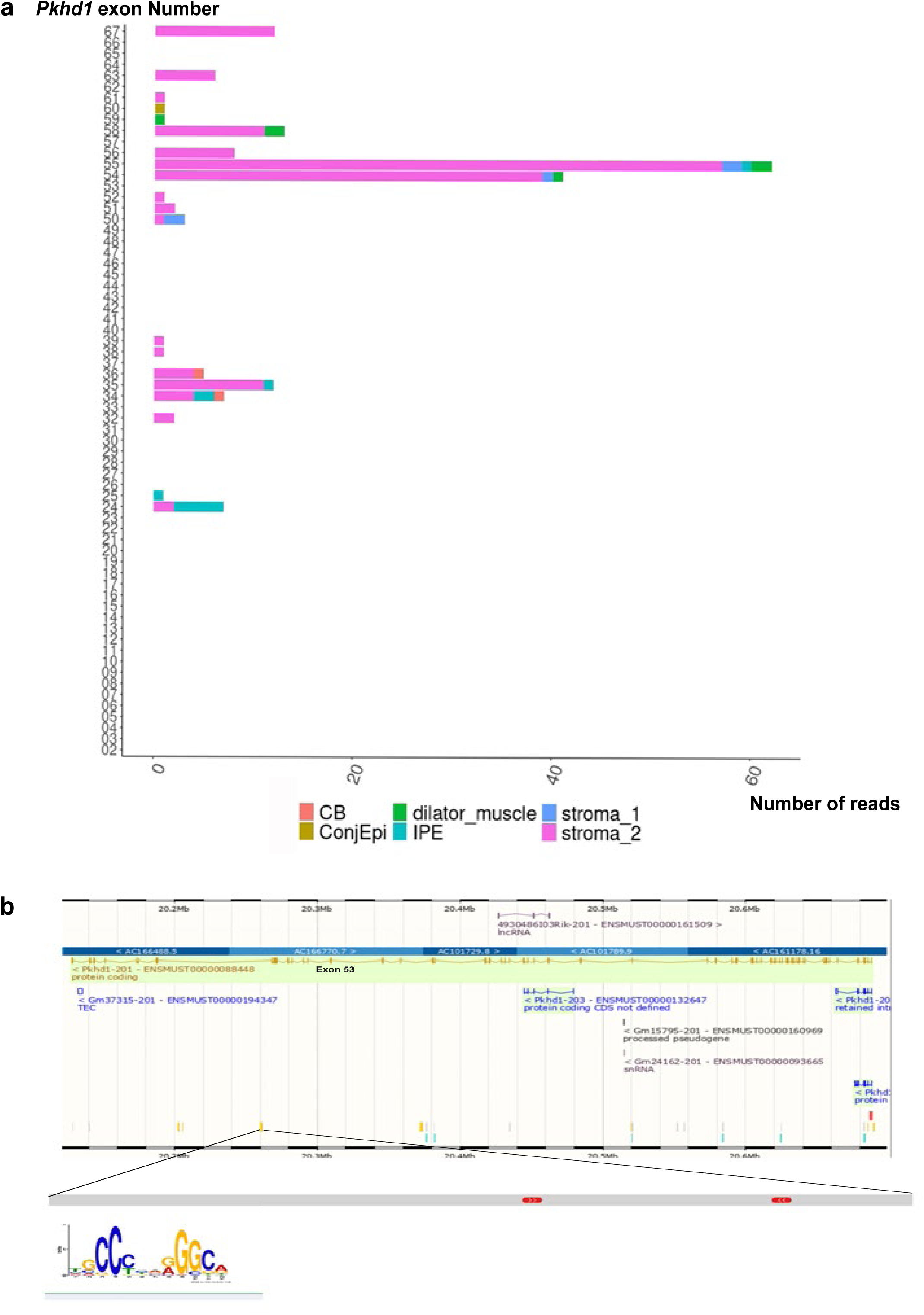
*Pkhd1* exon read number and potential enhancer region. (a) Number of reads found for each *Pkhd1* exon in select eye cell populations, colored by cell type. Note the high number of reads in genomic exons 54 and 55, particularly in stroma 2 cells. IPE: iris pigmented epithelium, CB: ciliary body, ConjEpi: conjunctival epithelium. (b) UCSC Genome Browser map of *Pkhd1* genomic region showing enhancers in yellow. An enhancer in the vicinity of exons 54 and 55 is enlarged to show the location of AP-2β motifs

**Supplementary Fig. 6:**
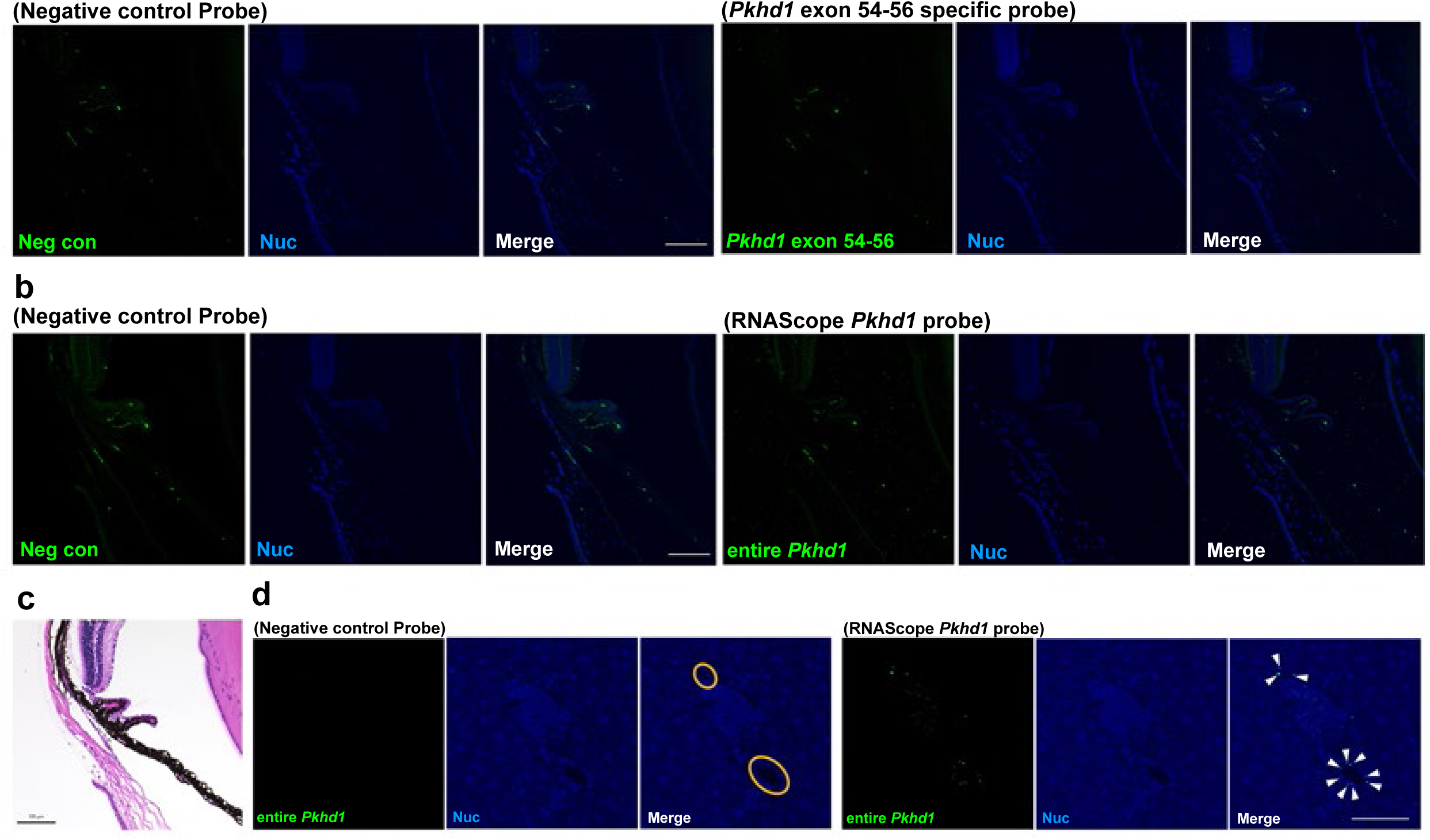
RNA *in situ* hybridization for *Pkhd1* transcripts in the eye. (a, b) Representative images of *in situ* hybridization of WT mouse eye at 2 months of age using RNAScope HiPlex v2. (a) Results for a negative control and a probe targeting *Pkhd1* exons 54-56 (Round 1). Scale bar 100 μm. The signal was similar in the two, consistent with a non-specific pattern. (b) Results for a negative control and a probe targeting the entire *Pkhd1* gene (Round 2). Scale bar 100 μm. No *Pkhd1*-specific signal was detected. (c) H&E staining of the sample used for *in situ* hybridization. Scale bar 100 μm. (d) Representative images of in situ hybridization of a wild-type mouse liver at 2 months of age using RNAScope HiPlex v2 probes for negative control and entire *Pkhd1* gene. Scale bar 50 μm. Upper: stained with negative control probe, bottom: stained with *Pkhd1* gene probe that targets the full-length transcript. Since *Pkhd1* is highly expressed in cholangiocytes, this experiment was conducted as a positive control for the eye study. Orange circles shown in the merged image of negative control probe indicate bile ducts. Arrowheads shown in the merged image of *Pkhd1* probe indicate *Pkhd1* transcripts localized to cholangiocytes. These results **indicate that the** *in situ* hybridization worked well but *Pkhd1* transcripts likely are below the level of detection of the assay in the eye.

**Supplementary Fig. 7:**
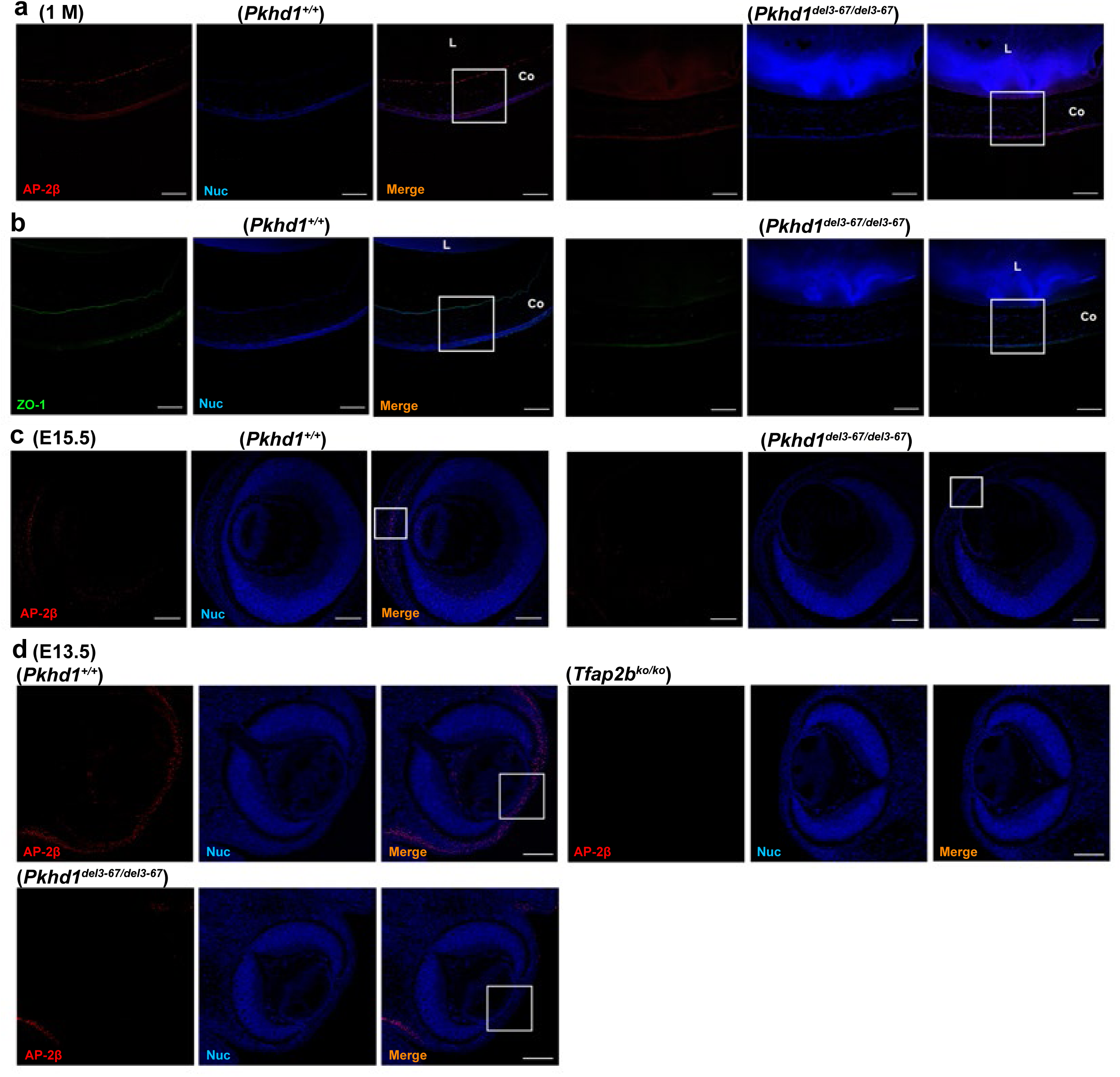
ZO-1 and AP-2β staining in *Pkhd1^del3-67/del3-67^* eyes. (a) Immunostaining of AP-2β in 1 month old WT and littermate *Pkhd1^del3-67/del3-67^* mouse eyes. AP-2β (red), and nuclear staining with Hoechst 33342 (blue). Scale bar 100 μm. The area highlighted in merged images is shown in Figure 3a. (b) Immunostaining of ZO-1 in specimens from the same mice in panel A. ZO-1 (green), and nuclear staining with Hoechst 33342 (blue). Scale bar 100 μm. The area highlighted in merged images is shown in Figure 3b. (c) Immunostaining of AP-2β in E15.5 WT and *Pkhd1^del3-67/del3-67^* mouse eyes. AP-2β (red), and nuclear staining with Hoechst 33342 (blue). Scale bar 100 μm. The area highlighted in merged images is shown in Figure 3c. (d) E13.5 control, *Tfap2b^ko/ko^* and *Pkhd1^del3-67/del3-67^* mouse eyes. AP-2β (red), and nuclear staining with Hoechst 33342 (blue). Scale bar 100 μm. The area highlighted in merged images is shown in Figure 3d. This image serves as a negative control for Figure 3d.

**Supplementary Fig. 8:**
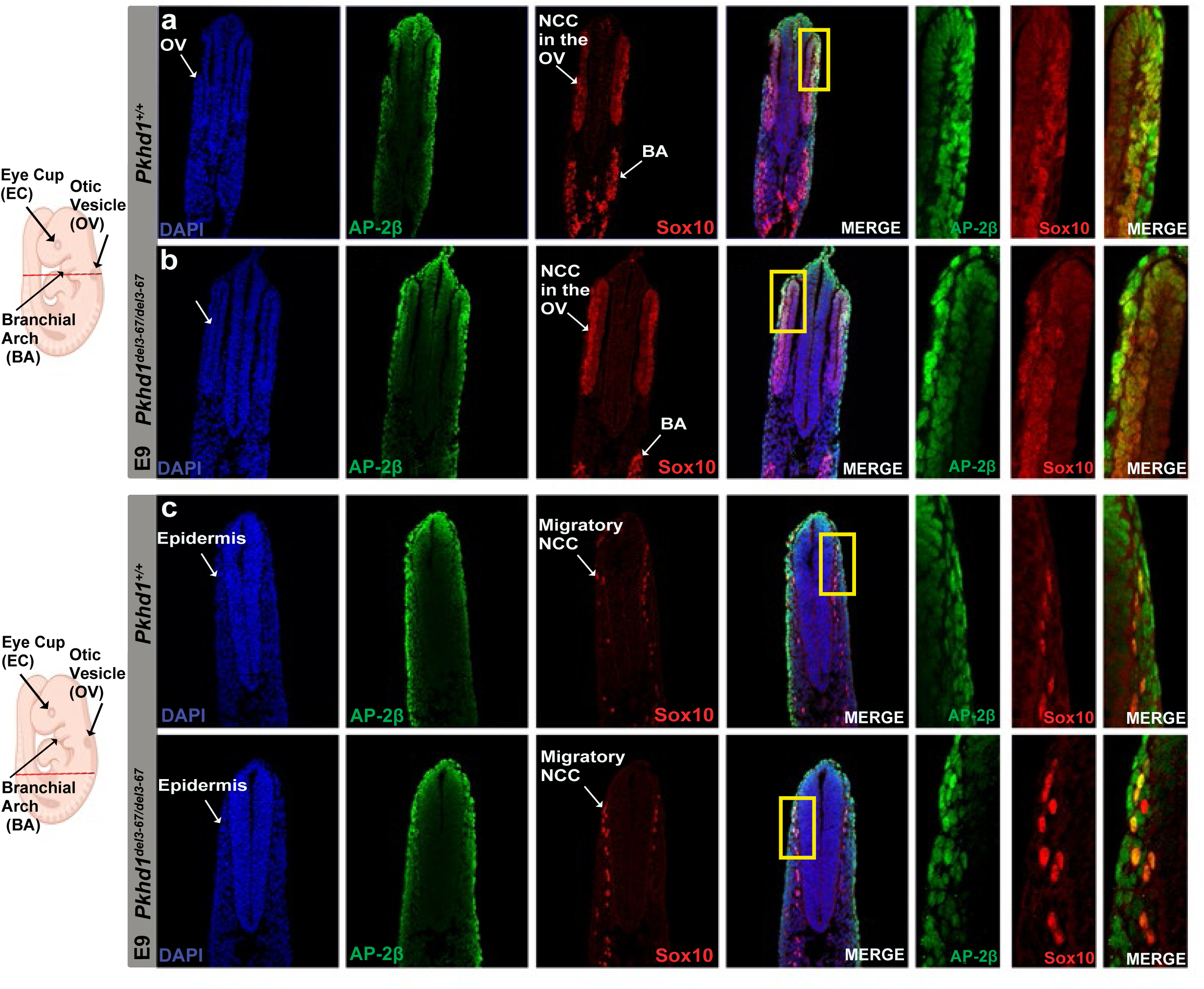
AP-2β expression in early embryonic mouse. On the left are two schematic diagrams of an E9 mouse embryo depicting with a red line the respective plane of cross section presented in Supplementary Figures 8a-d. (a, b) Cryosections from the hindbrain level showing equal levels of AP-2β (green) and Sox10 (red) expression in neural crest derived cells in the otic vesicle and branchial arches in wild type and mutant E9 embryos. (c, d) Cryosections from the posterior axial level show a similar pattern of migrating neural cells with equal levels of AP-2β (green) and Sox10 (red) expression in wild type and mutant E9 embryos. Yellow insets show a magnification of the squared area showing co-localization of AP-2β and Sox10 in the neural crest and its derivatives. Note that AP-2β is expressed also in the epidermis.

**Supplementary Fig. 9:**
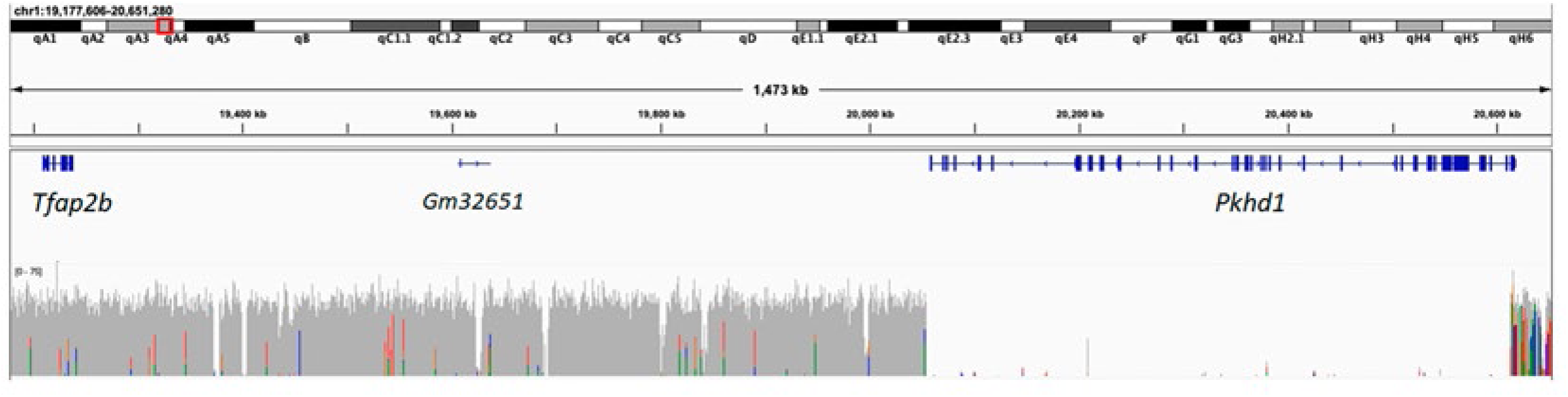
Structural variation analysis of Whole Genome Sequencing results of the *Pkhd1*^del3-67/del3-67^ mouse line. Whole genome sequence showing the *Pkhd1* and *Tfap2b* regions in a mutant *Pkhd1^del3-67/del3-67^* mouse, visualized using Integrative Genomics Viewer relative to mouse reference genome GRCm38/mm10. The vertical bars correspond to number of reads covering the region, color coded to show the proportion of reads with nucleotide mismatches (T: red; A: green; G: gold; C: blue) relative to the reference genome. Note the absence of reads in *Pkhd1* exons in the intended deleted region (chr1:20,053,820-20,613,713). Areas of low read density outside the deletion area correspond to genomic segments with high between-strain variability (Lilue, J., et al., D1243-D1249) or blacklisted as problematic sequencing regions (Amemiya, H., et al. *Sci Rep*, 2019. 9(1): p. 9354).

**Supplementary Fig. 10:**
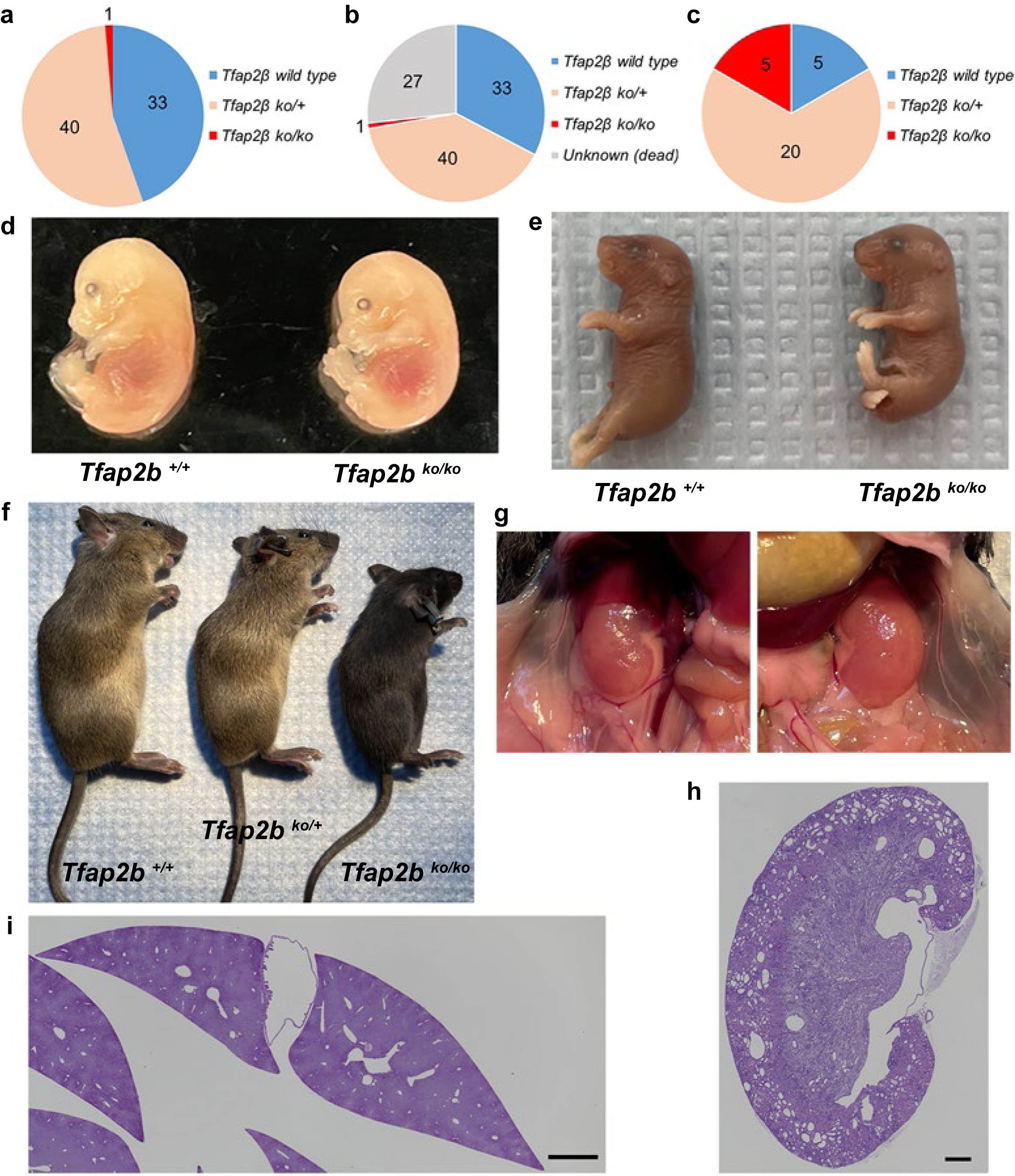
Additional data for *Tfap2b^ko/ko^* mouse. (a) Genotyping results of weaned pups from *Tfap2b^ko/+^* x *Tfap2b ^ko/+^* mating. Genotypes were determined for 74 pups weaned from 15 litters at 3 weeks of age. Only 1 *Tfap2b^ko/ko^* survived until weaning. (b) Genotyping results of delivered pups from *Tfap2b^ko/+^* x *Tfap2b ^ko/+^* mating. A total of 101 pups from 15 litters were delivered but 27 died before weaning and their genotypes could not be determined. Same cohort used in Supplementary Figure 10a. (c) Genotyping results of E 15.5 mouse embryos from *Tfap2b^ko/+^* x *Tfap2b ^ko/+^* mating. A total of 20 embryos from three litters were evaluated. The number of *Tfap2b^+/+^* and *Tfap2b^ko/ko^* embryos was the same at this age suggesting no early embryonic lethality in *Tfap2b^ko/ko^*. (d) Images of E13.5 *Tfap2b^+/+^* and *Tfap2b^ko/ko^* mouse embryo littermates. (e) Images of E18.5 *Tfap2b^+/+^* and *Tfap2b^ko/ko^* mouse embryo of littermates. (f) Images of P40 *Tfap2b^+/+^*, *Tfap2b^ko/+^* and *Tfap2b^ko/ko^* littermates. All mice are female. The sole surviving *Tfap2b^ko/ko^* was smaller than her littermates. (g) Images of the cystic kidneys of P40 *Tfap2b^ko/ko^* mouse. (h) Representative H&E staining of cystic kidney of *Tfap2b^ko/ko^*. Scale bar 500 μm. (i) Representative H&E staining of the liver of *Tfap2b^ko/ko^*. Scale bar 1000 μm. No cystic changes were observed in the liver of *Tfap2b^ko/ko^*.

**Supplementary Fig. 11.**
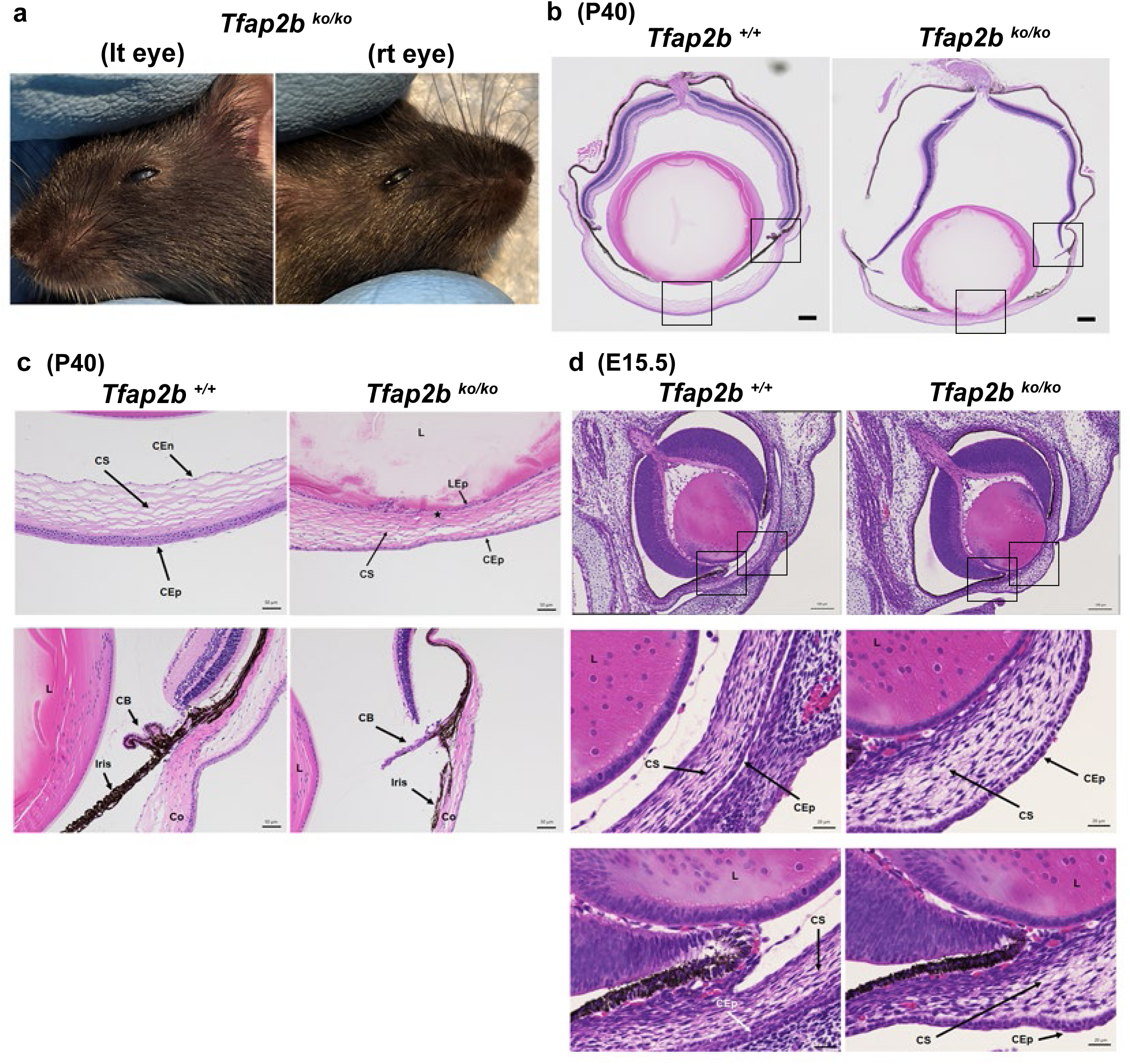
*Tfap2b^ko/ko^* mouse eye phenotype. (a) Image of the eyes of *Tfap2b^ko/ko^* at P40. The *Tfap2b^ko/ko^* mouse had developed corneal opacity in both eyes. (b) Representative eye pathology with H&E staining at P40. Whole eye images of P40 *Tfap2b^+/+^* and *Tfap2b^ko/ko^* littermates are shown. Black squares indicate the region shown in Supplementary Figure 11c. Scale bar 200 μm. (c) Representative eye pathology with H&E staining of P40 *Tfap2b^+/+^* and *Tfap2b^ko/ko^* littermates. Upper: cornea with 50 μm scale bar, bottom: angle tissue with 50 μm scale bar. CB: ciliary body, Co: cornea, CEn: corneal endothelium, CS: corneal stroma, CEp: corneal epithelium, L: lens, LEp: lens epithelium. star indicates absent corneal endothelium. (d) Representative eye pathology with H&E staining of E15.5 *Tfap2b^+/+^* and *Tfap2b^ko/ko^* littermate. Top: whole eye image with 100 μm scale bar. Black squares indicate the region of cornea and angle tissue. Middle: cornea with 20 μm scale bar, bottom: angle tissue with 20 μm scale bar. CS: corneal stroma, CEp: corneal epithelium, L: lens.

**Supplementary Fig. 12:**
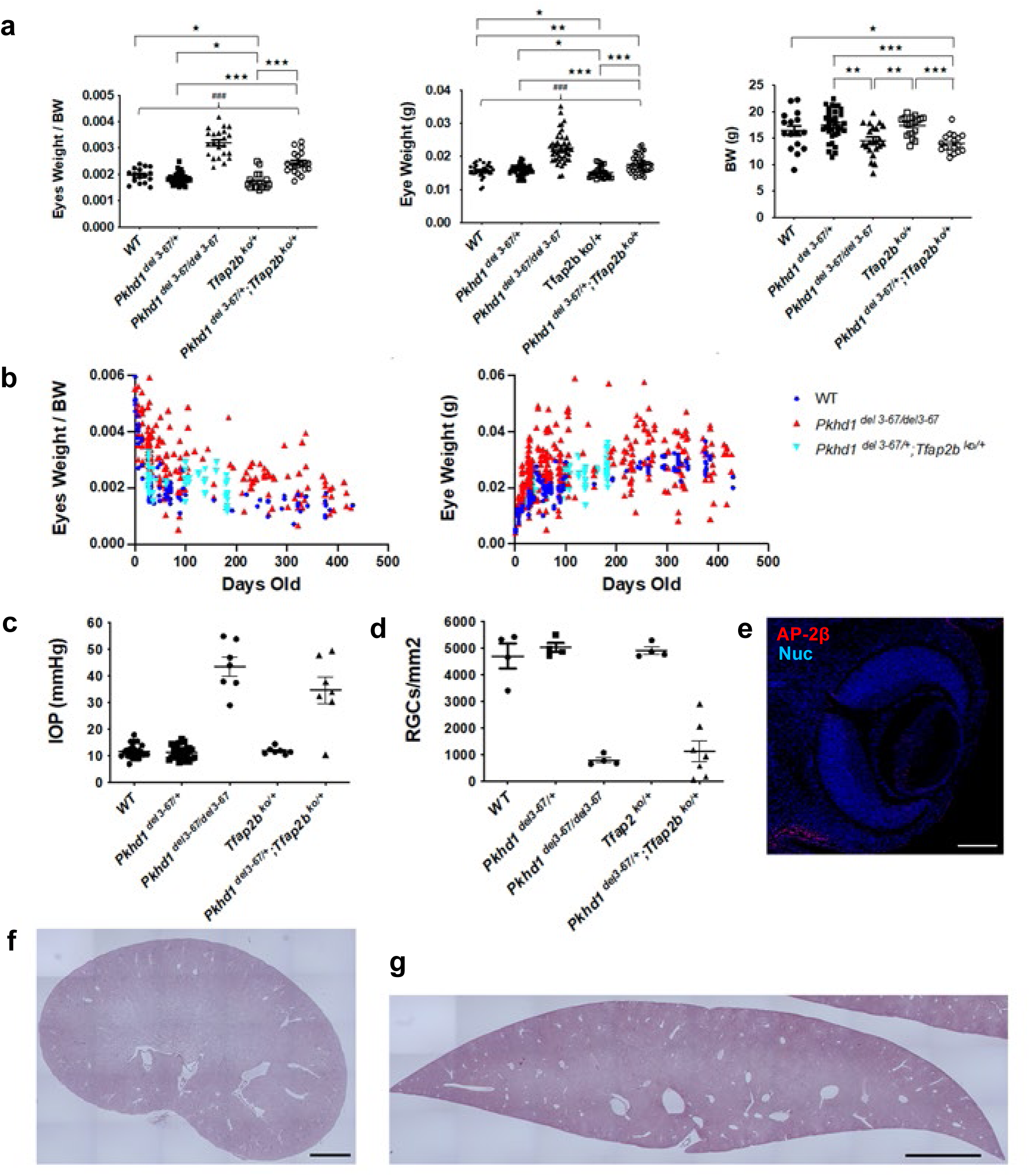
Additional data for *Pkhd1^del3-67+^;Tfap2b^ko/+^* trans-heterozygote mouse phenotype. (a) From left: Eyes weight per body weight (BW), eye weight and BW at 4 weeks of age including all mouse genotypes. A subset of these data is also presented in Figure 6e. N=16 mice (9 males and 7 females), *Pkhd1^del3-67+^* N=33 mice (15 males and 18 females), *Pkhd1^del3-67del3-67^* N=32 mice (16 males and 16 females), *Tfap2b^ko/+^* N=20 mice (16 males and 4 females), trans-heterozygote N=18 mice (7 males and 11 females). For Supplementary Figure 12 a and b, each dot in Eyes Weight/BW is the mean value of eyes from a single mouse while each dot in eye weight indicates the measurement for a single eye. Only significant differences are shown: ^★★★^ p < 0.001, ^★★^ p < 0.01. ^★^ P < 0.05. (b) Scatter plot of eyes weight per BW and eye weights of WT, *Pkhd1^del3-67/del3-67^* and *Pkhd1^del3-67+^;Tfap2b^ko/+^* trans-heterozygote at different ages. Other genotypes were not included to improve clarity. Left: scatter plots of eyes weight per BW. Right; eye weight (in grams). (c) IOP of all mouse genotypes at 3 weeks of age. A subset of these data is also presented in Figure 6f. Each dot is the mean value of IOP for a single mouse. (d) Number of RGC at 4 weeks of age from all mouse genotypes. A subset of these data is also presented in Figure 6g. Each dot represents the mean number of RGC counted from a series of immunostained images from a single mouse. (e) An E13.5 *Pkhd1^del3-67+^;Tfap2b^ko/+^* trans-heterozygote mouse eye stained for AP-2β showing absent staining in most structures but some positive cells in the POM, as had been seen with the E13.5 *Pkhd1^del3-67/del3-67^* specimens. This example illustrates the similar patterns of AP-2β staining observed in *Pkhd1^del3-67/del3-67^* and *Pkhd1^del3-67+^;Tfap2b^ko/+^* trans-heterozygous samples. Five of the *Pkhd1^del3-67+^;Tfap2b^ko/+^* trans-heterozygous eyes had the pattern shown in Figure 6l while one had the pattern shown here. Conversely, one E13.5 *Pkhd1^del3-67del3-67^* specimen lacked staining in the POM as shown in Figure 6f for the *Pkhd1^del3-67+^;Tfap2b^ko/+^*trans-heterozygote. AP-2β (red), and nuclear staining with Hoechst 33342 (blue). Scale bar 100 μm. (f) Representative H&E image of a non-cystic kidney of a *Pkhd1^del3-67+^;Tfap2b^ko/+^* trans-heterozygote. Scale bar 1000 μm. (g) Representative H&E image of the non-cystic liver of a *Pkhd1^del3-67+^;Tfap2b^ko/+^* trans-heterozygote. Scale bar 2000 μm.

**Supplementary Fig. 13:**
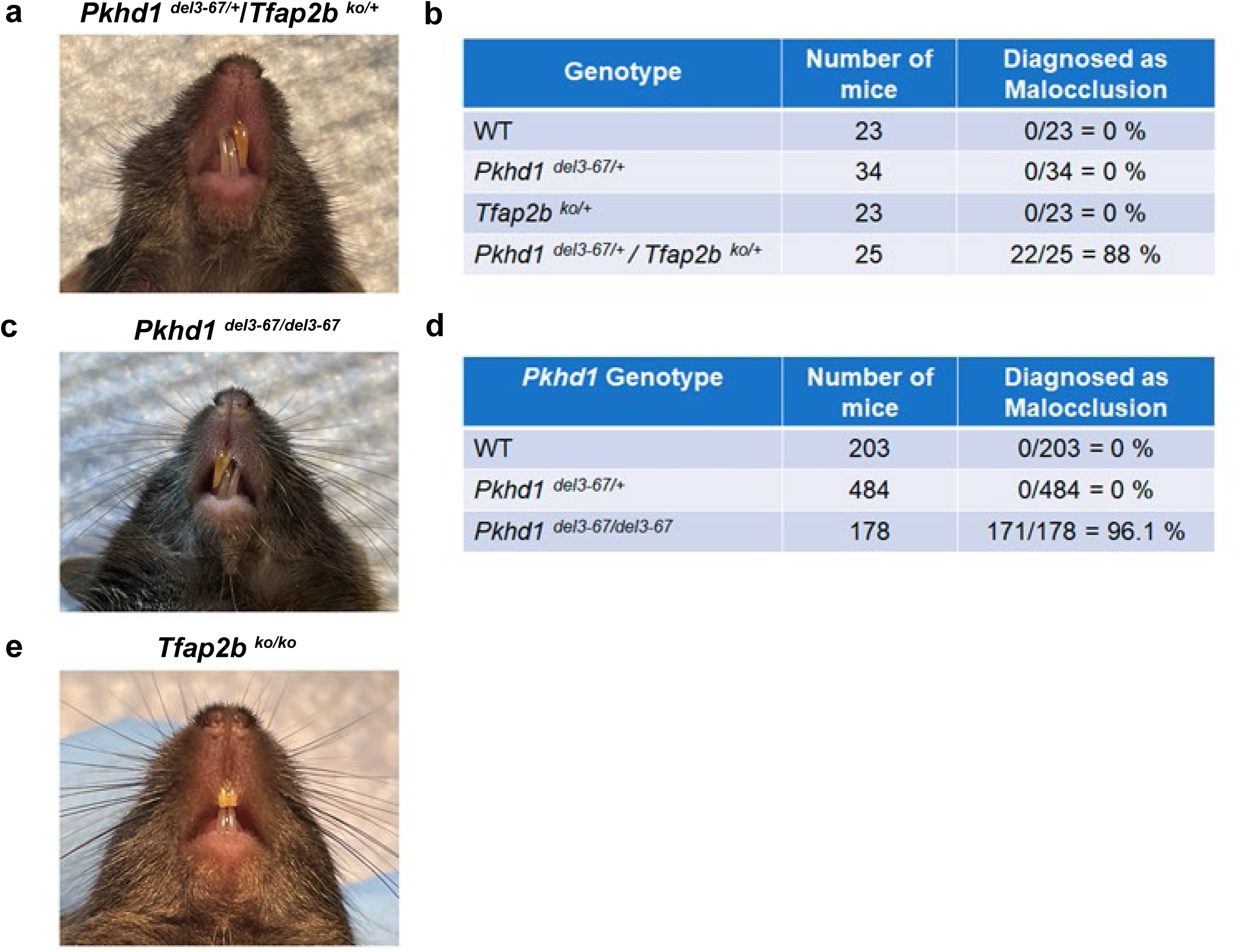
Incisor malocclusion phenotype. (a) Representative image of incisor malocclusion of a *Pkhd1^del3-67+^;Tfap2b^ko/+^* trans-heterozygous mouse. (b) Prevalence of malocclusion in pups related to the *Pkhd1^del3-67+^;Tfap2b^ko/+^* trans-heterozygous mouse line. Data include pups from *Pkhd1^del3-67/del3-67^* x *Tfap2b^ko/+^*mating in addition to the pups shown in Figure 6d. All mice that did not develop malocclusion developed eye phenotype (data not shown). (c) Representative image of incisor malocclusion of *Pkhd1^del3-67/del3-67^* mouse. (d) Prevalence of malocclusion in the *Pkhd1^del3-67^*mouse line. (e) Image of incisor of *Tfap2b^ko/ko^* mouse shown in Supplementary Figure 11a.

